# Increased pollination specialization did not increase corolla shape constraints in Antillean plants

**DOI:** 10.1101/041533

**Authors:** Simon Joly, François Lambert, Hermine Alexandre, Julien Clavel, Étienne Léveillé-Bourret, John L. Clark

## Abstract

Flowers show important structural variation as reproductive organs but the evolutionary forces underlying this diversity are still poorly understood. In animal-pollinated species, flower shape is strongly fashioned by selection imposed by pollinators, which is expected to vary according to guilds of effective pollinators. Using the Antillean subtribe Gesneriinae (Gesneriaceae), we tested the hypothesis that pollination specialists pollinated by one functional type of pollinator have maintained more similar corolla shapes through time due to stronger selection constraints compared to species with more generalist pollination strategies. Using geometric morphometrics and evolutionary models, we showed that the corolla of hummingbird specialists, bat specialists, and species with a mixed-pollination strategy (pollinated by hummingbirds and bats; thus a more generalist strategy) have distinct shapes and that these shapes have evolved under evolutionary constraints. However, we did not find support for smaller disparity in corolla shape for hummingbird specialists compared to more generalist species. This could be because the corolla shape of more generalist species in subtribe Gesneriinae, which has evolved multiple times, is finely adapted to be effectively pollinated by both bats and hummingbirds. These results suggest that pollination generalization is not necessarily associated with relaxed selection constraints.

## Introduction

The variation of flower shapes and structures we observe in nature is a constant reminder of the power of natural selection. This diversity is often attributed to zoophilous pollination, which has been associated with increased diversification in angiosperms (Stebbins, 1970; Crepet, 1984; Johnson, 2010; van der Niet and Johnson, 2012). Indeed, pollinator-driven selection pressure has been associated with species diversification (Whittall and Hodges, 2007) and frequent pollinator shifts often correlate with increased species diversification rates (e.g., Valente et al., 2012; Forest et al., 2014; Breitkopf et al., 2015). Yet, despite the numerous studies on pollination-driven selection at the population level (reviewed below), on the dissection of the genetic basis of several floral transitions between species pollinated by different pollinators (reviewed in: Galliot et al., 2006; Yuan et al., 2013) and of phylogenetic investigations of pollination systems at macroevolutionary levels (e.g., Perret et al., 2007; Smith, 2010), there is still a gap in our understanding on how the microevolutionary forces operating at the population level shape the macroevolutionary patterns we observe (Waser, 1998).

Selection can affect flower morphology differently when a population is adapting to a novel pollinator guild (transition phase) compared to when it is under the influence of a relatively constant pollinator guild (stasis phase). The transition phase is expected to involve strong directional selection until the population has a phenotype close to the optimum for the new pollinators (Lande, 1976). Studies on pollinator-mediated selection have found evidence for strong directional selection for flower shape in the transition phase (Galen, 1989), while others have shown that pollinators can drive flower colour transitions in populations (Waser and Price, 1981; Stanton et al., 1986). Although not a direct measurement of selection, the numerous studies reporting geographically structured flower variation associated with variation in pollinator guilds further support these findings (e.g., Gómez and Perfectti, 2010; Newman et al., 2014; Niet et al., 2014; Martén-Rodríguez et al., 2011), especially when reciprocal transplant experiments confirmed these patterns (Newman et al., 2012; Boberg et al., 2014; Sun et al., 2014).

For populations in stasis phase, that is with a relatively constant selection pressure from a stable pollinator guild, the floral traits are expected to be under stabilizing selection around optimal trait values. The mean phenotype of a population evolving under stabilizing selection is affected by both selection and drift, with selection pulling the mean phenotype towards the fitness optimum and drift due to finite population sizes moving it in random directions (Lande, 1976, 1979). Although stabilizing selection on floral traits have sometimes been observed in pollinator-mediated selection studies (Sahli and Conner, 2011; Conner et al., 2003), most studies failed to find such evidence (Campbell et al., 1991; O’Connell and Johnston, 1998; Maad, 2000). This might be because these phases are not so stable and that these studies are typically performed on a yearly basis. Indeed, studies have shown that selection on floral traits can vary from year to year in populations (Campbell, 1989; Campbell et al., 1991) due to temporal variation in pollinator abundance or environmental conditions. Nevertheless, there is considerable evidence that traits involved in the mechanical fit between the flower and the pollinators are under long-term stabilizing selection pressure as they show less variation in populations than other traits (Muchhala, 2006; Cresswell, 1998). Interestingly, these observations suggest that evidence for such stabilizing selection might be better studied over many generations, or even at macroevolutionary scales, than for a single generation (see also Haller and Hendry, 2014).

The intensity of constraints during the stasis phase is also expected to vary according to the level of pollination generalization of the species of interest. If the flower shape of specialist flowers should show evidence of stabilizing selection around an optimal shape adapted to its pollinator, the expectations are less clear for flowers of generalist species that possess pollinators that are functionally different for the plant (Aigner, 2001, 2006; Sahli and Conner, 2011). In general, unless the different functional pollinators all select for a common shape (common peak model: Sahli and Conner, 2011), generalists effectively pollinated by more than one functional type of pollinators are expected to be under weaker selection constraints than specialists (Johnson and Steiner, 2000). These predictions do not seem to have been tested thoroughly but are important to understand how and why flowers diversify under the selection of animal pollinators (Johnson, 2010).

In this study, we used a macroevolutionary approach to test whether increased specialisation in pollination strategies is associated with reduced corolla shape diversification (disparity) caused by stronger long-term selective constraints in species of the subtribe Gesneriinae of the Gesneriaceae family in the Caribbean islands. The recent development of powerful phylogenetic comparative methods allows the estimation of historic selective constraints on large groups of species (e.g., Hansen and Martins, 1996; Beaulieu et al., 2012; Butler and King, 2004) and thus testing specific hypotheses regarding the role of pollinators on floral trait evolution (e.g., Gómez et al., 2015). Unlike many investigations performed at the population level, such approaches aim at measuring selective constraints in terms of selective optima or in the rate at which disparity accrues over macroevolutionary scales and, as such, should be informative to understand the forces that have been determinant in modelling the morphology of large groups of species.

The subtribe Gesneriinae represents an ideal group to test this hypothesis. This diverse group in terms of floral morphologies is almost completely endemic to the Antilles and diversified into approximately 81 species (Skog, 2012) during the last 10 millions years (Roalson et al., 2008; Roalson and Roberts, 2016). The group has been the subject of several pollination studies that classified the species into different pollination syndromes that vary in their degree of ecological specialization (Martén-Rodríguez and Fenster, 2008; Martén-Rodríguez et al., 2009, 2010, 2015). There exists several definitions of pollination specialization/generalization, but globally plants pollinated by more species are considered more generalist (see papers in Waser and Ollerton, 2006), although information on the relative abundance (Medan et al., 2006) and functional diversity of pollinators (Johnson and Steiner, 2000; Fenster et al., 2004; Gómez and Zamora, 2006) should ideally taken into account. Here, we follow Fleming and Muchhala (2008) and measure ecological specialization with respect to the number of effective functional pollinator groups, with species pollinated by more functional pollinator groups being more generalists.

Specialist pollination strategies in Gesneriinae include hummingbird pollination, bat pollination, moth pollination and bee pollination (Fig. 1). Species with these strategies are pollinated by a single functional type (or guild) of pollinator and most often by a single species (Martén-Rodríguez and Fenster, 2008; Martén-Rodríguez et al., 2009, 2010, 2015). A fifth pollination strategy is considered more generalist as it is effectively pollinated in similar proportion by hummingbirds and bats (Martén-Rodríguez et al., 2009), two pollinators belonging to different functional groups that have different plant (growth form) and floral (nectar, shape, colour) preferences (Baker, 1961; Faegri and van der Pijl, 1979; Flemming et al., 2005). Although there exists many examples of more generalist species, these species are nevertheless ecologically more generalized than species pollinated by a single functional group of pollinators because they rely on more diversified resources (Gómez and Zamora, 2006). To avoid confusion with super-generalist species, we will use the term mixed– pollination strategy to refer to them in this study. Species of the Gesneriinae are sometimes visited by insects, but these always have marginal importance (Martén-Rodríguez and Fenster, 2008; Martén-Rodríguez et al., 2009, 2015) except for the insect pollination syndromes. A phylogenetic study of the group suggested multiple origins of most pollination strategies (Martén-Rodríguez et al., 2010), making it a perfect group to study selective forces acting on each one. In this study, we augmented previous phylogenetic hypotheses of the group by adding more species and genetic markers and we used geometric morphometrics of corolla shape and evolutionary models to test that (1) corolla shape evolution in the group supports distinct pollination syndromes, (2) corolla shape evolution is characterized by long-term constraints, and that (3) the corolla shape of pollination specialists show reduced disparity compared to the mixed-pollination species.

**Figure 1:**
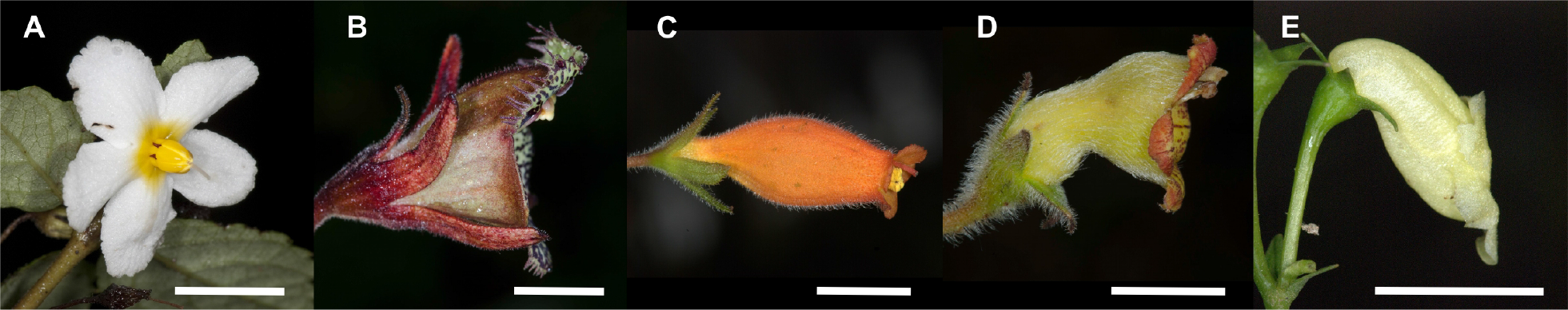
Gesneriinae flowers showing the different pollination strategies discussed in the study: (A) bee pollination (*Bellonia spinosa*, voucher: XXX); (B) bat pollination (*Gesneria fruticosa*, voucher: XXX); C) hummingbird pollination (*Rhytidophyllum rupincola*, voucher: XXX); D) mixed-pollination (*Rhytidophyllum auriculatum*, voucher: XXX); E) moth pollination (*Gesneria humilis*, voucher: XXX). The bar indicates 1 cm. Photographs by XXX.

## Material and Methods

### Floral morphology and pollination strategies

We took and collected photographs of 137 flowers in anthesis (137 distinct individuals, all from different localities) in longitudinal view, from 50 species (supplementary Table S1, S2; picture thumbnails are available as supplementary material) for a mean of 2.8 individuals per species (sd. dev. = 2.4). Most of these were taken in the wild, but a few specimens came from botanical gardens. We also took three pictures of the same flower (releasing and grabbing the pedicel between pictures) for four species at the Montreal Botanical Garden to quantify the error involved in hand-photographing the specimens as this is how most specimens were photographed.

Pollinator information was obtained from the literature (Martén-Rodríguez and Fenster, 2008; Martén-Rodríguez et al., 2009, 2010, 2015). Pollination strategy of species without field observation were inferred following the conclusions of Martén-Rodríguez et al. (2009). Briefly, hummingbird specialists have straight tubular corollas with bright colours and diurnal anthesis, bat specialists have green or white campanulate (bell-shaped) corollas with nocturnal anthesis and exserted anthers, and species with a mixed-pollination strategy are intermediate with subcampanulate corollas (bell-shaped with a basal constriction) showing various colours with frequent coloured spots, and diurnal as well as nocturnal anther dehiscence and nectar production (Martén-Rodríguez et al., 2009, Fig. 1). So far, only one moth pollinated species has been observed and it has a pale pouched corolla (Fig. 1). All analyses were performed (1) using only species with confirmed pollinator information and (2) also adding species with inferred strategies. We followed the taxonomy of Skog (2012) except for recent modifications in the *Gesneria viridiflora* complex (unpublished data).

### Molecular methods

A total of 94 specimens were included in the phylogenetic analyses (supplementary Table S3). *Koehleria* sp. ‘Trinidad’ (tribe Gesnerieae) and *Henckelia malayana* (tribe Trichosporeae) were included as outgroups. DNA was extracted using the plant DNA extraction kits from QIAGEN (Toronto, Ontario) or BioBasics (Markham, Ontario). Five nuclear genes were amplified and sequenced: *CYCLOIDEA*, *CHI*, *UF3GT*, *F3H*, *GAPDH*. The first four are unlinked (unpublished linkage map), whereas no data is available for *GAPDH*. Primer sequences and PCR conditions can be found in supplementary Table S4. Sequencing reactions were performed by the Genome Quebec Innovation Centre and run on a 3730xl DNA Analyzer (Applied Biosystems). Sequences from both primers were assembled into contigs and corrected manually in Geneious vers. 1.8. DNA sequences generated for this study were augmented with previously published sequences (supplementary Table S3).

### Phylogenetic analyses

Gene sequences were aligned using MAFFT (Katoh and Standley, 2013). Ambiguous alignment sections in intron regions of *CHI* and *GAPDH* were removed using gblocks (Castresana, 2000) with the default settings. Alignments were verified by eye and no obviously misaligned region remained after treatment with gblocks. Substitution models were selected by Akaike Information Criterion (AIC) with jModeltest 2 (Darriba et al., 2012) using an optimized maximum likelihood tree. A species tree was reconstructed using *BEAST in BEAST ver. 1.8.2 (Drummond et al., 2012). A Yule prior was chosen for the tree, a lognormal relaxed molecular clock for gene trees, and a gamma (2,1) prior for gene rates. Other parameters were left to the default settings. Three independent Markov Chain Monte Carlo (MCMC) analyses of 1 × 10^8^ generations were performed, sampling trees and parameters every 10,000 generations. Convergence of the runs was reached for parameter values, tree topology and clade posterior probabilities. The first 2 × 10^7^ generations were discarded as burnin and the remaining trees were combined for the analyses. The maximum clade credibility tree with median node heights was used for graphical representation.

### Geometric morphometric analyses

Six landmarks and 26 semi-landmarks were positioned on photographs using `tpsDig2` (Rohlf, 2010). Two landmarks were positioned at the base of the corolla, two at the tips of the petal lobes, and two at the base of the petal lobes, which generally corresponds to the corolla tube opening. The semi-landmarks were then positioned at equal distance along the curve of the corolla (13 on each side) between the landmarks at the base of the corolla and at the base of the petal lobes. The sepals were present on most of the pictures. The landmark data was imported in `R` (R core team, 2014) where it was transformed by generalized Procrustes analysis using the `geomorph R` package (Adams and Otárola-Castillo, 2013). The semi-landmarks on curves were slid along their tangent directions during the superimposition by minimizing the Procrustes distance between the reference and target specimen (Bookstein, 1997). Size was not considered in the analyses because we were interested in shape and because a scale was not available for all specimens. Because the actinomorphic flowers of bee pollinated species (*Bellonia ssp.*) do not allow homologous placement of landmarks, these were dropped from the morphometric analyses.

Landmarks were positioned twice for each photograph and a Procrustes ANOVA quantified the variance explained by these technical replicates, which were combined for the remaining analyses. We also used a Procrustes ANOVA to quantify the variation among the replicated photographs of the same flowers; these replicates were not included in the final analyses. The Procrustes aligned specimens were projected into the tangent space, hereafter the morphospace, using Principal Component Analysis (PCA) of the covariance matrix using the `prcomp` function in `R`.

To characterize the total morphological variation for each pollination strategy, we estimated the distance of the mean corolla shape of each species to the pollinator strategy centroid in multivariate space and tested if these distances were different for the different pollination strategies using the `betadisper` function of the `vegan` package in `R` (Oksanen et al., 2017). The differences were tested by ANOVA. We also partitioned the variation into intraspecific and interspecific components for each pollination strategy using Procrustes ANOVA, reporting adjusted *R*^*2*^ values.

Morphological integration (Klingenberg, 2013) was quantified using the variance of the eigenvalues of a PCA on the covariance matrix (Pavlicev et al., 2009; Klingenberg, 2013), scaling the eigenvalues by the total variance of the sample to get an index independent of the total sample variation (Young, 2006). This was estimated on all individuals for the hummingbird and mixed-pollination species. Bat specialists were omitted from this analysis because there were too few species to give a result comparable to the other pollination strategies.

### Ancestral states reconstruction

Ancestral state reconstruction was performed to estimate the probability of all pollination strategy states for all nodes of the phylogeny. The best transition model was first selected by second order AIC (AICc) with the `geiger R` package (Harmon et al., 2008). Eight models selected based on biological relevance were compared. The Equal Rate (ER), Symmetric (SYM) and All Rates Different (ARD) were tested with modified versions that give a single rate to and from the moth and bee states (ER.2, SYM.2, and ARD.2). In addition, a 4-rate model was tested where rates differed according to the actual state and a single rate to and from the bee and moth states, and finally a 3-rate model with one rate for transitions from and to bee and moth states, one from hummingbirds to bats or mixed-pollination, and a third from bat or mixed-pollination to all states except bee or moth. Using the best model, the joint ancestral state probabilities were estimated using stochastic character mapping (Huelsenbeck et al., 2003) on the maximum clade credibility tree with 2000 simulated character histories. When estimating ancestral states with only species with confirmed pollinators, the other species were given equal prior probabilities in the simulations. To estimate the number of transitions between states while accounting for phylogenetic uncertainty, 500 character histories were simulated on 2000 species trees randomly sampled from the posterior distribution from the species tree search using the `phytools R` package. The median number of transitions between all states from all simulated character histories were reported as well as 95% credible intervals.

### Evolutionary constraints on flower shape

Given the nature of the hypotheses tested, two types of evolutionary models based on the Brownian motion (BM) and the Ornstein-Uhlenbeck (OU) stochastic processes were considered. BM models the accumulation of independent and infinitesimal stochastic phenotypic changes (controlled by the drift rate parameter *σ*^2^) along the branches of a phylogeny; it can approximate various scenarios of phenotypic evolution such as drift, fluctuating directional selection or punctuated change (Felsenstein, 1985; Hansen and Martins, 1996; O’Meara et al., 2006). In contrast, the OU process models selection toward a common optimal trait value (Felsenstein, 1988; Hansen and Martins, 1996) and adds to the BM model a selection parameter *α* that determines the strength of selection towards an optimal trait *θ* (details on the models can be found in Hansen and Martins, 1996; Butler and King, 2004; Beaulieu et al., 2012). When the strength of selection is null (*α* = 0), the OU process reduces to BM. These models can be made more complex, for instance by allowing parameters to vary in different parts of the tree (selective regimes - e.g., Butler and King, 2004; O’Meara et al., 2006; Beaulieu et al., 2012) and are therefore useful for characterizing the evolutionary constraints of the pollination strategies.

Generalist pollination is hypothesized to promote phenotypic diversification (disparity) of corolla shape because it is thought to be under weaker selection (Johnson and Steiner, 2000), but also because of the spatio-temporal variation in pollinator abundance that could result in fluctuating selection pressures (Herrera, 1988) or in a variety of species- or population-specific adaptive peaks (see Discussion). As such, mixed-pollination species are expected to best fit a BM process. In contrast, due to their adaptation to a single functional pollinator, pollination specialists are expected to show smaller variation around a better defined shape optimum and thus fit an OU process. However, BM and OU processes can be difficult to distinguish, and an OU process can best fit the data for other reasons such as measurement error (Silvestro et al., 2015), bounded trait variation (Boucher and Démery, 2016) or small sample sizes (Cooper et al., 2016). In contrast, OU models are less likely to be selected when analyzing the primary axes of variation from a PCA (Uyeda et al., 2015). Therefore, prediction of higher phenotypic disparity is often better assessed through evaluation of parameters estimated under OU or BM for species pollinated by different functional groups of pollinators. For instance, with the BM process, the drift rate (*σ*^2^) describes the accumulation of phenotypic variance over the tree and is therefore tightly related to phenotypic disparity (O’Meara et al., 2006; Thomas et al., 2006; Price et al., 2013). Following our hypothesis of lower phenotypic disparity for pollination specialists, we predict they should have a smaller *σ*^2^ compared to mixed-pollination species. Similarly, under an OU model, the stationary variance around an optimum, expressed as *σ*^2^/2*α* for the univariate case, is also tightly related to phenotypic disparity. We thus expect pollination specialists to be associated with stronger corolla shape constraints (i.e., higher *α* values) and smaller stationary variances compared to mixed-pollination species in models where either *α* or *σ*^2^ vary between strategies (see below). Finally, we expect phenotypic evolutionary correlations between traits inferred from multivariate comparative models to be higher in pollination specialists (i.e., higher phenotypic integration, see for instance Revell and Collar, 2009).

We evaluated and compared the model fit and parameter estimates with the predictions of our hypotheses using univariate and multivariate models because they allow investigating different aspects of the data. Univariate models allowed us to fit a greater range of evolutionary models that are not yet implemented in multivariate approaches and allow investigating if different shape components evolved under similar constraints. In contrast, multivariate models allow to fit an evolutionary model on several shape components at once and also allow to investigate patterns of evolutionary correlation among traits for the different pollination strategies; that is, studying phenotypic integration in an evolutionary context. For univariate models, we fitted BM models with one drift rate for the whole tree (BM1) and with one rate per regime (BMV), but also versions that allow different ancestral states for the different regimes (O’Meara et al., 2006; Thomas et al., 2009); model BM1m has distinct trait means per regime but a single drift rate across the tree, while BMVm has distinct means and drift rates for each regime. We also fitted different variants of the OU models (Beaulieu et al., 2012): with a single optimum *θ* (OU1), with different optima for lineages with different pollination strategies (OUM), different *θ* and selective strength *α* (OUMA), different *θ* and rates of stochastic motion *σ*^2^ (OUMV), or different *θ*, *α* and *σ*^2^ (model OUMVA) for the different pollination strategies. We also considered ecological release models, in which one regime on the tree is evolving under BM and the other under an OU process, either with a shared drift rate *σ*^2^ (OUBM and BMOU) or with their own drift rates (models OUBMi and BMOUi) which are sometimes called ecological release and radiate models (see Slater, 2013). The model OUBM considers hummingbird specialists to be evolving under an OU model whereas the mixed-pollination species are evolving under a BM model, and vice versa. Several multivariate models were also considered: BM1, BMV, BM1m, BMVm, OU1, OUM, OUBM, BMOU, OUBMi, and BMOUi. The multivariate OU models allowing different con-traints on different regimes (OUMA, OUMV, OUMVA) are not implemented yet and thus we can not estimate regime specific evolutionary covariance (or correlation) matrices. However, we expect such models to be over-parameterized with respect to the number of species considered in our study.

In the remaining, we therefore consider the comparison of phenotypic evolutionary correlations obtained from the *σ*^2^ correlation matrices of the multivariate BM models only. Yet, focussing on the interpretation of parameters obtained under the BM processes can be misleading if BM is a poor descriptor of the phenotypic evolution (see for instance Price et al., 2013). To make sure this did not affect our estimates, we simulated datasets using a OUM model on 100 trees randomly selected from the posterior distribution using the parameters estimated from the observed data. We then fitted these simulated data with the BMVm model to obtain *σ*^2^ correlation matrices that were compared with the original *σ*^2^ correlation matrices.

The models were fitted for the first three principal components of the morphospace using the `R` packages `mvMORPH` (Clavel et al., 2015) and `OUwie` (Beaulieu et al., 2012). The models were fitted on a sample of 1000 trees from the posterior distribution of species trees on which the character history was inferred by one instance of stochastic mapping (Huelsenbeck et al., 2003) using maximum likelihood in the `phytools R` package (Revell, 2012). This accounts for phylogenetic uncertainty and the stochasticity of the character state reconstructions (Revell, 2013). All the trees were re-scaled to unit height. Intraspecific variation was taken into account by using the sampling variance (the squared standard error) of species as measurement error in model fitting; species without biological replicates were given the mean squared standard error of species with the same pollination strategy. The models were compared using *AICc* weights that can be roughly considered as the relative weight of evidence in favour of a model given a set of models (Burnham and Anderson, 2002). The analyses were performed with inferred pollination strategies as well as with species with confirmed pollination strategies only. Note that because there were few confirmed bat pollinated species and a single moth pollinated species, species with these pollination strategies were excluded from the analyses. However, the inclusion of bat pollinated species in the univariate models did not affect the conclusions (data not shown). The data and scripts used to replicate all analyses are available as supplementary information.

## Results

### Phylogeny

The species phylogeny showed that the bee pollinated genus *Bellonia* is sister to the rest of the subtribe, and the subtribe (*Bellonia* + *Gesneria* + *Rhytidophyllum*) received a posterior probability of 1 (not shown). *Rhytidophyllum* and *Gesneria* were found to form distinct clades, although *Gesneria* received weaker support (Fig. 2). This reinforces the distinction between these two genera, which has been debated over the years. There is one exception, *Rhytidophyllum bicolor*, which is included for the first time in a molecular phylogeny and that falls within *Gesneria*. The taxonomic name of this species will have to be reconsidered. Several branches show strong clade posterior probabilities, but some had less support due to lack of phylogenetic signal or conflict between gene trees, indicating the importance of incorporating phylogenetic uncertainty in the following analyses.

**Figure 2:**
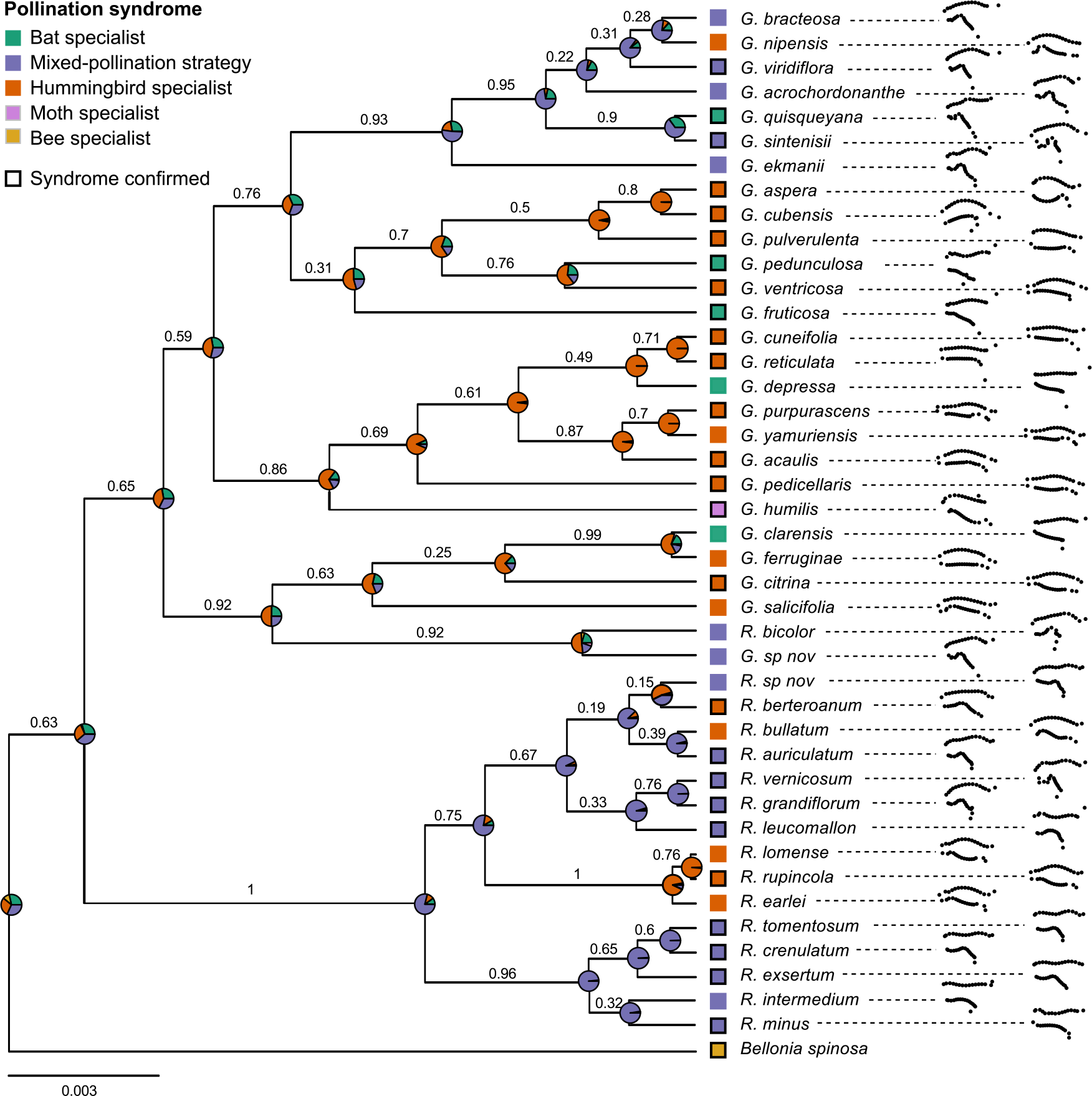
Species phylogeny showing mean corolla shapes (after Procrustes analysis). Pollination strategies are shown with those that have been confirmed indicated by a black contour. Pie charts represent the joint probability of each state at nodes as estimated by stochastic mapping from only species with confirmed pollinators. Clade posterior probabilities are shown above branches. Outgroup taxa are not shown.

The best character evolutionary model (smallest *AICc*) was the 3 rates model with one rate for transitions from and to the bee and moth states, one from hummingbirds to bats or mixed-pollination, and a third from bat or mixed-pollination to all states except bee and moth. Ancestral state reconstruction (Fig. 2) suggests that the hummingbird pollination is the most likely ancestral state for the *Gesneria* clade, although it is only slightly more likely than an ancestral mixed-pollination strategy. In contrast, the mixed-pollination strategy is the most probable ancestral state for the *Rhytidophyllum* clade. A hummingbird pollinated ancestor for the subtribe is more probable, but only very slightly. This reflects the difficulty in estimating the ancestral states for nodes near the root of a phylogeny (Gascuel and Steel, 2014). The ancestral state reconstruction with the inferred pollination strategies (Fig. S1) were highly similar to those of Fig. 2.

Estimation of the number of transitions supports several transitions between the bat, the mixed-pollination and the hummingbird strategies (Table 1). The number of transition from mixed-pollination to hummingbird and from mixed-pollination to bat was slightly higher than from bat to mixed-pollination and bat to hummingbird, which was also slightly higher than the number of transitions from hummingbird to bats and hummingbird to mixed-pollination (Table 1). However, because the confidence intervals largely overlap, we can conclude that the number of transitions between these three main pollination strategies are not significantly different. The results were almost identical when analyses were performed with inferred pollination strategies (Supplementary Table S5). These estimates are similar to those of Martén-Rodríguez et al. (2010), although they found fewer reversals to hummingbirds in their study. Overall, these results confirm multiple evolutionary origins for all pollination strategies except for the bee and moth (95 % CI always > 2; Table 1).

**Table 1:**
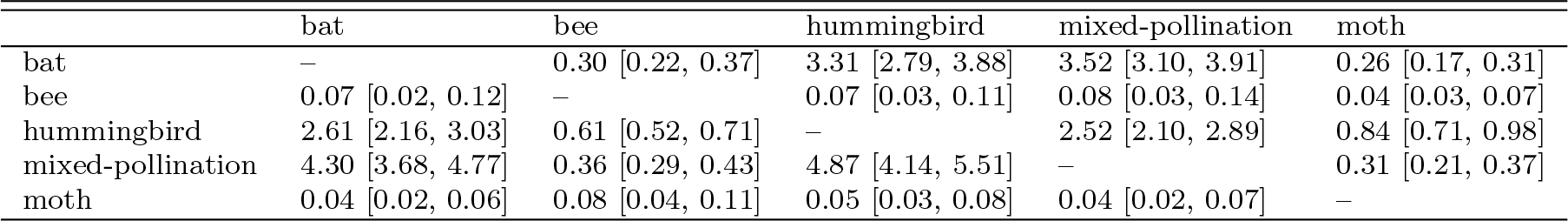
Number of transitions between the different pollination strategies according to stochastic mapping. The median values obtained from the character simulations over the posterior distribution of species tree is reported as well as 95% credible intervals. Ancestral states are in rows.

|  | bat | bee | hummingbird | mixed-pollination | moth |
| --- | --- | --- | --- | --- | --- |
| bat | – | 0.30 [0.22, 0.37] | 3.31 [2.79, 3.88] | 3.52 [3.10, 3.91] | 0.26 [0.17, 0.31] |
| bee | 0.07 [0.02, 0.12] | – | 0.07 [0.03, 0.11] | 0.08 [0.03, 0.14] | 0.04 [0.03, 0.07] |
| hummingbird | 2.61 [2.16, 3.03] | 0.61 [0.52, 0.71] | – | 2.52 [2.10, 2.89] | 0.84 [0.71, 0.98] |
| mixed-pollination | 4.30 [3.68, 4.77] | 0.36 [0.29, 0.43] | 4.87 [4.14, 5.51] | – | 0.31 [0.21, 0.37] |
| moth | 0.04 [0.02, 0.06] | 0.08 [0.04, 0.11] | 0.05 [0.03, 0.08] | 0.04 [0.02, 0.07] | – |

### Corolla shape

We found only 0.15% of variation between independent pictures of the same flower in the replication experiment, which is lower than the variation involved in the landmark positioning (0.81%). Therefore, we conclude that the error included in the data by the picture acquisition was minimal. Similarly, because the technical replicates accounted for only 0.56% of the total variance in the final dataset, the mean shape between replicates was used for the remaining analyses.

The morphospace explained 79% of the total shape variance in the first three axes. The first principal component (PC) represents 53.6% of the variance and is characterized by campanulate vs. tubular corollas (Fig. 3A), broadly differentiating hummingbird specialists from the other species. This concurs with a previous study that showed that this was indeed the main characteristic differentiating the hummingbird pollination strategy from the bat and the mixed-pollination strategies (Martén-Rodríguez et al., 2009). PC2 explains 20.6% of the variance and is characterized by corolla curvature and distinguished the moth pollinated *G. humilis*. The bat and the mixed-pollination strategies could not be differentiated with this PCA, but a second PCA that excluded moth and hummingbird pollinated species (both confirmed and inferred) found that the bat and mixed-pollination strategies were separated along PC3 that is characterized by a basal constriction in the corolla (Fig. 3B), a character known to distinguish bat pollinated species (that generally lack the constriction) from species with a mixed-pollination strategy (Martén-Rodríguez et al., 2009). The single bat pollinated species that groups with mixed-pollination species on this axis is *Gesneria quisqueyana* (see interactive supplementary figures for information on the individual and species positioning in the PCAs), which, in contrast to other bat pollinated species in the group, excludes hummingbirds during the day by actively closing its flowers (Martén-Rodríguez et al., 2009).

**Figure 3:**
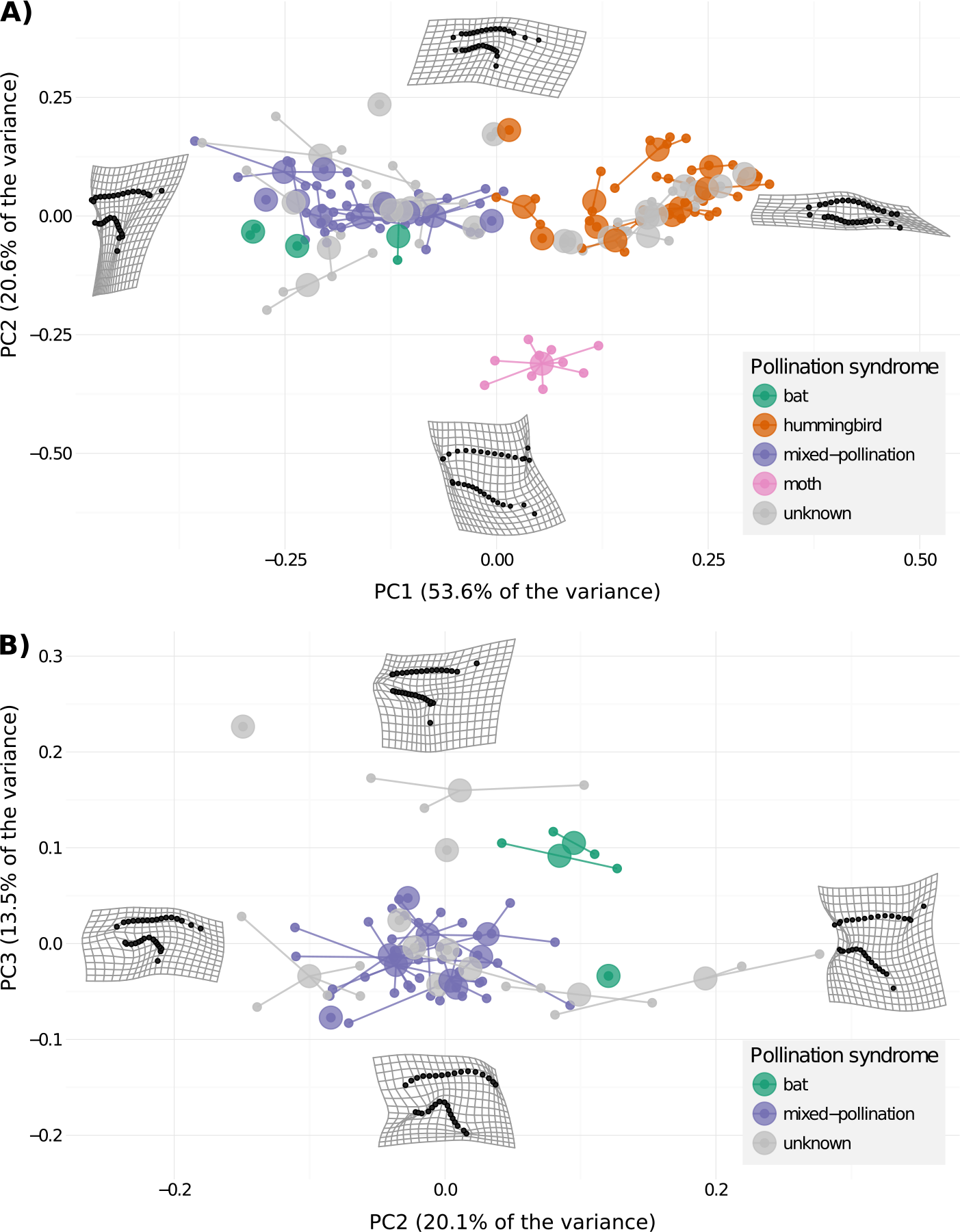
Principal component analyses showing the corolla shape morphospace for (A) all species and (B) species excluding hummingbird (both confirmed and inferred) and moth pollinated species. The large dots on the plot represent the species means, which are connected by a line to the floral shapes of the individuals belonging to the species (small dots). Thin-plate spline deformation grids show corolla shape variation along the principal components (plus or minus 2 standard deviation from the mean shape). *Bellonia spinosa* (bee pollinated) was not included in the morphometric analyses because it has a radial symmetry.

### Variation partitioning

The pollination strategies did not have a significantly different corolla variation among species (ANOVA: F = 1.92, df = 2, *p* = 0.1654). The partitioning of the shape variance for the different pollination strategies showed that the proportion of variance explained among species corresponded to 81.4% (*p* < 0.001) for hummingbird pollinated species, 91.3% (*p* = 0.22) for bat pollinated species and 50.4% (*p* < 0.001) for mixed-pollination species. The result of the variance partitioning for the bat pollinated species should be interpreted with caution because there was only three species with less than two replicated individuals on average within species for this syndrome.

### Morphological integration

Flower components are generally well integrated as they develop, function and evolve jointly (Ashman and Majetic, 2006), a concept called morphological integration (reviewed in Klingenberg, 2013). A large morphological integration index supports important integration because morphological variation is concentrated in few principal components. The results showed that species with a mixed-pollination strategy had a slightly greater morphological integration (0.0069) than hummingbird pollinated species (0.0050).

### Evolutionary models

#### Univariate models

For PC1 that captures variation in corolla opening, all models that received *AICc* weights greater than zero suggest that the hummingbird specialists and the mixed-pollination species differed in their mean shape as they all included distinct *θ* for the two strategies (Table 2). The best models, OUM and BM1m (*AICc* weight of 0.48 and 0.35, respectively), suggest that the two pollination strategies had similar evolutionary phenotypic variance as they constrain them to have identical parameters. This trend is also supported by parameter estimates of supported models that allowed the strategies to differ in drift rates (BMVm) or stationary variance (OUMV, OUMA, OUMVA) as these estimates were very similar for the two strategies (Table 2). The phylogenetic half-time of the OUM model, which corresponds to the time required for the expected phenotype to move half-way towards the optimal shape from its ancestral state (Hansen, 1997), was of 0.009. Given that the trees were scaled to unit height, this small value imply either very strong selective pressure or a lack of phylogenetic correlation. The results of the analyses that included species with inferred pollination strategies were almost identical in terms of model selection and phenotypic disparity (Table S6).

**Table 2:** Parameter values of the univariate evolutionary models fitted on the first three principal components of the morphospace when only the species with confirmed pollinators were included in the analyses. Mean values from the posterior distribution of species trees are given for the AICc weights, whereas median values are given for the parameter estimates. Numbers in brackets indicate the 25% and the 75% quantiles. The best model for each PC is in bold. The *θ* parameter indicate the global or regime means (ancestral states) for the BM-type and OUBM-type models, whereas it indicates the stationary optimum trait for the OU-type models, *station*_*hum*_ and *station*_*mix*_ are the stationary distributions of the hummingbird and mixed-pollination strategies.

| PC1 |  |  |  |  |  |  |  |  |  |  |
| --- | --- | --- | --- | --- | --- | --- | --- | --- | --- | --- |
| Models | p | AICc weight | $\theta_{hum}$ | $\theta_{mix}$ | $\sigma_{hum}^2$ | $\sigma_{mix}^2$ | $station_{hum}$ | $station_{mix}$ | | |
| BM1 | 2 | 0 [0,0] | 0.057 [0.05,0.066] | 0.057 [0.05,0.066] | 0.046 [0.037,0.074] | 0.046 [0.037,0.074] | — | — |  |  |
| BMV | 3 | 0 [0,0] | 0.07 [0.011,0.113] | 0.07 [0.011,0.113] | 0.035 [0.02,0.093] | 0.045 [0.026,0.086] | — | — |  |  |
| BM1m | 3 | 0.35 [0.02,0.68] | 0.15 [0.144,0.155] | -0.123 [-0.131,-0.112] | 0.017 [0.013,0.023] | 0.017 [0.013,0.023] | — | — |  |  |
| BMVm | 4 | 0.1 [0.01,0.17] | 0.146 [0.138,0.153] | -0.124 [-0.132,-0.112] | 0.021 [0.014,0.031] | 0.013 [0.012,0.016] | — | — |  |  |
| OU1 | 3 | 0 [0,0] | 0.05 [0.039,0.057] | 0.05 [0.039,0.057] | 0.088 [0.061,0.197] | 0.088 [0.061,0.197] | 0.029 [0.029,0.031] | 0.029 [0.029,0.031] |  |  |
| OUM | 4 | 0.48 [0.11,0.84] | 0.16 [0.152,0.163] | -0.155 [-0.156,-0.152] | 0.859 [0.265,3.004] | 0.859 [0.265,3.004] | 0.006 [0.006,0.006] | 0.006 [0.006,0.006] |  |  |
| OUMV | 5 | 0.05 [0,0] | 0.197 [0.18,0.222] | -0.134 [-0.226,-0.094] | 8.248 [3.772,23.45] | 7.442 [2.802,21.646] | 2.684 [1.381,5.179] | 2.696 [1.443,4.635] |  |  |
| OUMA | 5 | 0.02 [0,0] | 0.195 [0.178,0.23] | -0.14 [-0.246,-0.09] | 8.935 [3.876,19.47] | 8.935 [3.876,19.47] | 2.573 [1.857,3.386] | 2.667 [2.107,3.739] |  |  |
| OUMVA | 6 | 0.01 [0,0] | 0.195 [0.175,0.212] | -0.141 [-0.257,-0.106] | 7.756 [3.12,23.395] | 6.743 [2.518,18.899] | 2.439 [1.388,4.782] | 3.018 [1.516,4.646] |  |  |
| OUBMi | 4 | 0 [0,0] | 0.125 [0.106,0.142] | 0.125 [0.106,0.142] | 0.252 [0.065,4.186] | 0.042 [0.023,0.064] | 0.009 [0.007,0.013] | — |  |  |
| BMOUi | 4 | 0 [0,0] | 0.038 [-0.03,0.092] | 0.038 [-0.03,0.092] | 0.04 [0.021,0.095] | 0.06 [0.032,0.104] | — | 460558.818 [0.017,1824288] |  |  |
| OUBM | 3 | 0 [0,0] | 0.135 [0.12,0.151] | 0.135 [0.12,0.151] | 0.073 [0.056,0.13] | 0.073 [0.056,0.13] | 0.007 [0.006,0.009] | — |  |  |
| BMOU | 3 | 0 [0,0] | 0.043 [-0.045,0.059] | 0.043 [-0.045,0.059] | 0.053 [0.037,0.107] | 0.053 [0.037,0.107] | — | 18437.122 [0.009,55497877] |  |  |
| PC2 |  |  |  |  |  |  |  |  |  |  |
| Models | p | AICc weight | $\theta_{hum}$ | $\theta_{mix}$ | $\sigma_{hum}^2$ | $\sigma_{mix}^2$ | $station_{hum}$ | $station_{mix}$ | | |
| BM1 | 2 | 0 [0,0] | -0.035 [-0.037,-0.032] | -0.035 [-0.037,-0.032] | 0.021 [0.013,0.038] | 0.021 [0.013,0.038] | — | — |  |  |
| BMV | 3 | 0.04 [0,0.04] | -0.035 [-0.041,-0.025] | -0.035 [-0.041,-0.025] | 0.036 [0.021,0.065] | 0 [0,0] | — | — |  |  |
| BM1m | 3 | 0 [0,0] | -0.031 [-0.035,-0.028] | -0.041 [-0.048,-0.036] | 0.021 [0.013,0.038] | 0.021 [0.013,0.038] | — | — |  |  |
| BMVm | 4 | 0.01 [0,0.01] | -0.028 [-0.036,-0.02] | -0.038 [-0.046,-0.026] | 0.036 [0.02,0.064] | 0 [0,0] | — | — |  |  |
| OU1 | 3 | 0.01 [0,0.02] | -0.036 [-0.036,-0.036] | -0.036 [-0.036,-0.036] | 0.972 [0.18,1.104] | 0.972 [0.18,1.104] | 0.003 [0.003,0.003] | 0.003 [0.003,0.003] |  |  |
| OUM | 4 | 0 [0,0.01] | -0.042 [-0.044,-0.042] | -0.027 [-0.027,-0.026] | 0.499 [0.107,1.041] | 0.499 [0.107,1.041] | 0.003 [0.003,0.003] | 0.003 [0.003,0.003] |  |  |
| OUMV | 5 | 0.15 [0.01,0.28] | -0.026 [-0.039,-0.014] | -0.017 [-0.04,-0.01] | 10.239 [5.932,18.219] | 0.018 [0,0.072] | 1.663 [1.314,2.512] | 0.003 [0,0.01] |  |  |
| OUMA | 5 | 0 [0,0] | -0.024 [-0.04,-0.015] | -0.016 [-0.038,-0.005] | 6.319 [3.531,11.251] | 6.319 [3.531,11.251] | 0.868 [0.671,1.145] | 0.868 [0.671,1.145] |  |  |
| OUMVA | 6 | 0.02 [0,0.03] | -0.025 [-0.038,-0.013] | -0.017 [-0.041,-0.011] | 13.96 [6.714,26.595] | 0.027 [0.001,1.106] | 1.772 [1.316,3.017] | 0.004 [0,0.167] |  |  |
| OUBMi | 4 | 0.72 [0.53,0.93] | -0.043 [-0.045,-0.041] | -0.043 [-0.045,-0.041] | 0.305 [0.132,0.93] | 0 [0,0] | 0.004 [0.004,0.004] | — |  |  |
| BMOUi | 4 | 0.01 [0,0.01] | -0.035 [-0.041,-0.025] | -0.035 [-0.041,-0.025] | 0.036 [0.021,0.065] | 0.001 [0,0.001] | — | 0 [0,0] |  |  |
| OUBM | 3 | 0 [0,0] | -0.044 [-0.05,-0.035] | -0.044 [-0.05,-0.035] | 0.03 [0.019,0.064] | 0.03 [0.019,0.064] | 0.003 [0.003,0.005] | — |  |  |
| BMOU | 3 | 0.03 [0,0.02] | -0.029 [-0.038,-0.025] | -0.029 [-0.038,-0.025] | 0.036 [0.021,0.064] | 0.036 [0.021,0.064] | — | 0 [0,0] |  |  |
| PC3 |  |  |  |  |  |  |  |  |  |  |
| Models | p | AICc weight | $\theta_{hum}$ | $\theta_{mix}$ | $\sigma_{hum}^2$ | $\sigma_{mix}^2$ | $station_{hum}$ | $station_{mix}$ | | |
| BM1 | 2 | 0.06 [0.01,0.09] | 0.017 [0.015,0.019] | 0.017 [0.015,0.019] | 0.005 [0.004,0.006] | 0.005 [0.004,0.006] | — | — |  |  |
| BMV | 3 | 0.02 [0,0.03] | 0.017 [0.015,0.019] | 0.017 [0.015,0.019] | 0.005 [0.004,0.006] | 0.005 [0.004,0.006] | — | — |  |  |
| BM1m | 3 | 0.02 [0,0.03] | 0.023 [0.022,0.026] | 0.004 [0.001,0.007] | 0.005 [0.004,0.005] | 0.005 [0.004,0.005] | — | — |  |  |
| BMVm | 4 | 0.01 [0,0.01] | 0.024 [0.022,0.026] | 0.004 [0.001,0.007] | 0.005 [0.004,0.006] | 0.005 [0.004,0.006] | — | — |  |  |
| OU1 | 3 | 0.18 [0.04,0.27] | 0.013 [0.013,0.016] | 0.013 [0.013,0.016] | 0.036 [0.017,0.739] | 0.036 [0.017,0.739] | 0.002 [0.002,0.002] | 0.002 [0.002,0.002] |  |  |
| OUM | 4 | 0.05 [0.01,0.07] | 0.015 [0.014,0.019] | 0.012 [0.011,0.013] | 0.03 [0.017,0.713] | 0.03 [0.017,0.713] | 0.002 [0.002,0.002] | 0.002 [0.002,0.002] |  |  |
| OUMV | 5 | 0.36 [0.18,0.53] | 0.027 [0.025,0.029] | 0.014 [0.01,0.022] | 13.351 [7.115,26.325] | 3.692 [2.121,6.126] | 1.033 [0.744,1.621] | 0.291 [0.215,0.359] |  |  |
| OUMA | 5 | 0.12 [0.04,0.17] | 0.026 [0.022,0.028] | 0.017 [0.01,0.024] | 4.885 [3.11,13.095] | 4.885 [3.11,13.095] | 0.505 [0.395,0.728] | 0.505 [0.395,0.728] |  |  |
| OUMVA | 6 | 0.04 [0.02,0.06] | 0.027 [0.024,0.029] | 0.014 [0.009,0.022] | 6.412 [4.403,19.508] | 2.492 [1.504,5.957] | 0.734 [0.556,1.073] | 0.264 [0.196,0.375] |  |  |
| OUBMi | 4 | 0.01 [0,0.01] | 0.015 [0.013,0.018] | 0.015 [0.013,0.018] | 0.012 [0.009,0.021] | 0.005 [0.004,0.006] | 0.002 [0.002,0.003] | — |  |  |
| BMOUi | 4 | 0.04 [0.01,0.04] | 0.015 [0.014,0.017] | 0.015 [0.014,0.017] | 0.005 [0.004,0.006] | 0.041 [0.021,0.233] | — | 0.001 [0.001,0.001] |  |  |
| OUBM | 3 | 0.03 [0,0.04] | 0.017 [0.014,0.02] | 0.017 [0.014,0.02] | 0.007 [0.006,0.009] | 0.007 [0.006,0.009] | 0.003 [0.002,0.004] | — |  |  |
| BMOU | 3 | 0.07 [0.01,0.07] | 0.015 [0.013,0.017] | 0.015 [0.013,0.017] | 0.006 [0.005,0.008] | 0.006 [0.005,0.008] | — | 0.001 [0.001,0.001] |  |  |

The PC2 of the morphospace that represents variation in the curve of the corolla was found to best fit a OUBMi model (*AICc* weight = 0.72; Table 2), with the hummingbird pollinated species evolving under a OU model and the mixed-pollination species evolving under a BM model, each with their own drift rate implying that this model cannot be simply interpreted as reduced constraints for mixed-pollination species. Nevertheless, the model suggests that the pollination strategies have the same mean shape for PC2 and that the two pollination strategies have evolved under different types of constraints. The median phylogenetic halftime was of 0.02 for the hummingbird species, suggesting either very strong selective pressure or a lack of phylogenetic correlation. Parameter estimates for the other models, in particular the second best model OUMV (*AICc* weight = 0.15), also supported similar mean shapes for the two pollination strategies and suggest that hummingbird pollinated species have greater phenotypic disparity as they have a greater stationary variance than mixed-pollination species (Table 2). The median phylogenetic half-time for the OUMV model was estimated to be 0.23, suggesting moderate constraints on corolla shape. The analyses including species with inferred pollination strategies best supported a OU1 model (*AICc* weights = 0.69; Table S6) indicating a lack of evidence for different constraints or disparity for the two strategies. But in models where the evolutionary rate of stationary variance was allowed to vary between strategies, the hummingbird pollinated species showed higher variance than the mixed-pollinated species (Table S6).

The PC3 that represents variation in the reflexion of the petal lobes was found to best fit a OUMV model (*AICc* weight = 0.36), although models OU1 and OUMA also received considerable weights (*AICc* weights of 0.18 and 0.12, respectively; Table 2). All three models suggest that this shape component tend to stay closer to the evolutionary mean than would be expected under a BM model. The OU1 suggests that the pollination strategies have the same mean shape, whereas the OUM and OUMV models suggest different mean shapes, although parameter estimates for these later models showed that the mean shapes for both strategies are not very far from each other (Table 2). The models OUMV and OUMA suggest different shape disparity with the hummingbird specialists having a higher stationary variance than mixed-pollination species. The models OUMV, OU1 and OUMA all suggested strong constraints with estimated phylogenetic half-times of 0.11, 0.08, and 0.14, respectively. In analyses with species with inferred pollination strategies, the OU1 model received the highest weight (0.30), although several models received weights greater than 0.05 (Table S6). As for the analyses with only species with confirmed pollination strategies, the hummingbird pollinated species showed higher stationary variance in models in which this parameter was allowed to vary between strategies (Table S6).

In some instances, the models OU1 and OUM did not always converge to the maximum likelihood solution when fitted with `OUwie`, especially for PC1. This is why we always fitted these models with `mvMORPH`, which is also faster. Similarly, the models OUMV, OUMA, and OUMVA showed relatively poor convergence and should be interpreted with caution.

#### Multivariate models

The multivariate analyses supported OUM as the best fitting model (*AICc* weight = 0.60; Table 3). This model suggests that the shape components have different evolutionary means for the two pollination strategies and that there is an evolutionary force that maintains the corolla shape closer to this evolutionary mean than would be expected under a BM model. The shape means estimated under the multivariate OUM model for each PC were very similar to that of the univariate estimates, as were the estimates of the stationary variance and phylogenetic halftimes (compare Tables 2 and 4). The stationary variance estimates were also similar to the observed variance among species for hummingbird pollinated species (PC1: 0.0068, PC2: 0.0049, PC3: 0.0041) and mixed-pollinated species (PC1: 0.0075, PC2: 0.0014, PC3: 0.0016), suggesting that the model is very close to be stationary.

**Table 3:** Model performance with the multivariate evolutionary models fitted on the first three principal components of the morphospace when only confirmed species are included in the analyses. The mean values obtained from the posterior distribution of species trees are given; numbers in brackets indicate the 25% and the 75% quantiles. The best model is in bold.

| Models | $\log Lik$ | Parameters | $AICc$ weight |
| --- | --- | --- | --- |
| BM1 | 67.98 [63.43,76.25] | 9 | 0.00 [0.00,0.00] |
| BMV | 80.44 [76.94,86.18] | 15 | 0.00 [0.00,0.00] |
| BM1m | 78.52 [72.7,86.95] | 12 | 0.24 [0.00,0.45] |
| BMVm | 90.47 [85.96,96.59] | 18 | 0.13 [0.00,0.12] |
| OU1 | 82.30 [74.47,87.04] | 15 | 0.02 [0.00,0.00] |
| **OUM** | **96.24 [93.47,98.18]** | **18** | **0.60 [0.03,1.00]** |
| OUBM | 77.17 [74.14,82.47] | 15 | 0.00 [0.00,0.00] |
| BMOU | 80.47 [76.97,85.85] | 15 | 0.01 [0.00,0.00] |
| OUBMi | 95.24 [93.99,97.05] | 21 | 0.01 [0.00,0.00] |
| BMOUi | 83.71 [80.01,89.11] | 21 | 0.00 [0.00,0.00] |

Because the current implementation do not allow the estimation of regime-specific evolutionary correlations between traits under the multivariate OUM model, we looked at the evolutionary correlations between drift rates (*σ*^2^) under the BMVm model, which was the third best supported model (*AICc* weight = 0.13; Table 3), to estimate the morphological integration for the two pollination strategies. The evolutionary correlations between shape components were always greater for the mixed-pollination strategy in terms of absolute correlation, although there is some uncertainty in these estimates as evident from the 50% posterior intervals (Fig. 4). Furthermore, the better support for the OUM and BM1m models also suggests that differences between pollination strategies are probably marginal or that we lack statistical power to detect significant differences. Because these correlations were obtained on a BMVm model whereas a OUM model was the one that received the highest support, there is a risk that the younger mixed-pollination clades may appear to have evolved faster under the BMVm model (Price et al., 2013), which could in turn affect the observed correlations. However, this does not seem to be the case as the correlations estimated on data simulated with the OUM model were similar between pollination strategies (Fig. 4), rejecting the possibility that the greater absolute correlations observed for the mixed-pollination strategy were due to model mis-specification. The multivariate results obtained when species with inferred pollinators were included were similar, with even more support for the OUM model (*AICc* weight = 1; Table S8, S9). However, the correlation between traits suggest phenotypic integration of more similar amplitude for the two pollination strategies with inferred pollinators (Fig. S2); these differences could be due to the small sample sizes as such correlations are difficult to estimate accurately.

**Figure 4:**
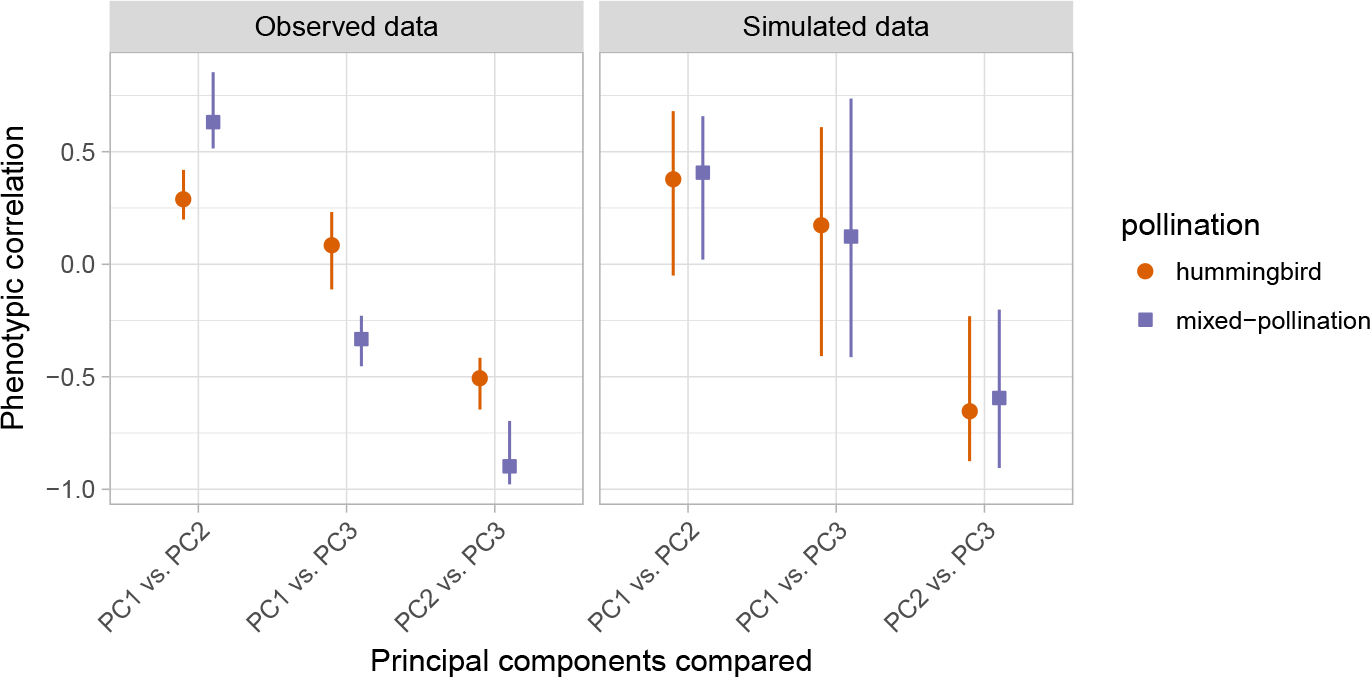
Graphical representation of the evolutionary correlations (i.e., standardized evolutionary rates matrices) obtained with the BMVm multivariate model with only species with confirmed pollination strategies, for the observed data (left panel) and for data simulated under the best fitting model (OUM; right panel). Symbols represent the median correlation and the lines the 25% and 75% quantiles for both hummingbirds and mixed-pollination strategies. No artifactual differences are detected between the two groups when fitting models on traits simulated with the OUM model and thus with a common evolutionary covariance (right panel, see text).

## Discussion

Although many aspects of the flower are required for assuring successful reproduction, the corolla shape is critical for the adaptation of plants to pollinators. In many species, the corolla guides the pollinator to allow precise pollen deposition on its body (Muchhala, 2007). But pollinators can also show an inherent preference for some floral shapes (Gómez et al., 2008) and can associate shape and reward when these are correlated (Meléndez-Ackerman et al., 1997). Floral shape has in fact repeatedly been shown to be under selection in pollination-driven selection studies (Galen, 1989; Campbell et al., 1991; O’Connell and Johnston, 1998; Maad, 2000) and can be sufficient to impose adaptive trade-off between pollinators (Muchhala, 2007). Even the corolla shape of super generalist species has been shown to adapt to particular guilds of pollinators (Gómez and Perfectti, 2010; Gómez et al., 2015).

In the Antillean genera *Gesneria* and *Rhytidophyllum*, pollination syndromes are well characterized and have good predictive value (Martén-Rodríguez et al., 2009), but previous studies were based on attractive and mechanical floral characters. Our results based on geometric morphometrics showed that it is possible to distinguish corollas of hummingbird pollinated species from moth pollinated species, and, although to a lesser degree, the corolla shapes of species with bat or mixed-pollination strategies. These conclusions were reinforced by the strong support in favour of distinct shapes for hummingbird specialists and mixed-pollination species in evolutionary models, both based on parameter estimates and on support for models supporting different evolutionary shape means (BMm models) or distinct shape optima (OUM models). These results, in addition to the fact that each pollination strategy evolved repeatedly in the Gesneriinae, further support the concept of pollination syndromes in this group and underlines the importance of corolla shape in floral adaptation to pollinators.

### Long-term evolutionary constraints on corolla shape

If several studies estimated selection pressures on flowers within populations, few have addressed the question at macroevolutionary scales (but see Gómez et al., 2015). We hypothesized, based on theoretical expectations and empirical results, that the corolla shape of flowers should have evolved under evolutionary contraints to maintain effective pollination, thereby reducing the phenotypic variance among species, and that this variance should be smaller for more specialist species because they are potentially under a more constant and precise selective pressure.

All analyses performed, both univariate and multivariate and using only species with confirmed pollinator information or also including species with inferred strategies, selected OU models that possess a *α* parameter that maintains the corolla shape closer to an evolutionary optimum. This supports the hypothesis that the corolla shape in the group has been affected by long-term evolutionary constraints, which could be interpreted as a consequence of the selective pressure imposed by pollinators. However, the analyses found very small phylogenetic halftimes, which are suggestive of very strong selection pressures and/or lack of phylogenetic correlation in the data (note that a strong impact of selection will necessarily reduce the phylogenetic correlation in the data). Considering a potential origin of the group ca. 10 mya (Roalson et al., 2008; Roalson and Roberts, 2016) and taking the smallest phylogenetic halftime obtained (0.002, for the PC1 in the multivariate analysis; Table 4), this means that a corolla shape can move half-way to its optimal shape in 0.002 × 10 = 0.02 million years, or 20,000 years. This is rapid, but not impossible considering that transitions between pollination strategies are generally driven by few genes of major effects (Galliot et al., 2006; Yuan et al., 2013). These small phylogenetic halftimes can also imply a lack of phylogenetic influence, but given that the corolla shape is clearly under genetic control in the group (Alexandre et al., 2015), this seems less plausible.

**Table 4:** Model parameters for the multivariate OUM model, which was the model that received the highest AICc weight (Table 3). The mean values obtained from the posterior distribution of species trees are given; numbers in brackets indicate the 25% and the 75% quantiles. The complete stationary variance matrix is given in Table S7.

| parameters | PC1 | PC2 | PC3 |
| --- | --- | --- | --- |
| $\theta_{hum}$ | 0.161 [0.152,0.166] | -0.043 [-0.046,-0.042] | 0.013 [0.009,0.015] |
| $\theta_{mix}$ | -0.156 [-0.159,-0.154] | -0.026 [-0.027,-0.023] | 0.013 [0.012,0.02] |
| $\sigma^2$ | 1.198 [0.135,0.135] | 1.328 [0.184,0.184] | 0.757 [0.005,0.005] |
| Phylogenetic halftime | 0.002 [0.001,0.003] | 0.01 [0.003,0.031] | 0.101 [0.01,0.194] |
| Stationary variance | 0.006 [0.005,0.006] | 0.003 [0.003,0.003] | 0.002 [0.002,0.002] |

Contrarily to our initial hypothesis, we did not find evidence that pollination specialist species show reduced phenotypic disparity compared to mixed-pollination species. The non-phylogenetic approaches suggested similar amount of variation among species for both pollination strategies, and this pattern was confirmed by the evolutionary models. Indeed, almost all analyses selected a model in which both strategies evolved under shared constraints, but for different means for each selective regime. Moreover, although the differences were marginal, the parameter estimates of the evolutionary models that allows the two strategies to have different phenotypic disparities almost constantly indicated that it was the hummingbird specialists that showed a higher disparity compared to the more generalist mixed-pollination species.

Morphological integration and evolutionary correlations between shape components allows us to take another view at evolutionary contraints on corolla shape. Indeed, important integration between the shape components suggests tight coordination for proper functioning and strong evolutionary correlations suggest that components have evolved in an highly coordinated fashion. The results showed both higher morphological integration and evolutionary correlations for the mixed-pollination species, which goes against the generally accepted idea that more generalist species are less constrained. Overall, we come to the conclusion that greater generalization in pollination strategies did not imply a relaxation of evolutionary constraints over macroevolutionary scales in Antillean Gesneriinae.

How can we explain that we did not observe more phenotypic disparity for the mixed-pollination strategy? Although any answer to this question is necessarily tentative at this point, a potential line of enquiry could be associated with the process by which the species become generalized. One such process, which we call the compromised phenotype and that motivated our initial hypotheses, is that generalists evolve by becoming intermediate in morphology relative to specialist species (Fig. 5). This could occur in the presence of weak or non-symmetric trade-off effects if both pollinators are present (Aigner, 2001, 2006; Sahli and Conner, 2011). Under such a scenario, the floral variation of generalists could span the whole region between the shapes of the two specialists either because of drift or because of variation in the the relative contribution of the different pollinators for each species. This could result in a relatively broad variation among species compared to that of specialists; but this is not what we observe here.

**Figure 5:**
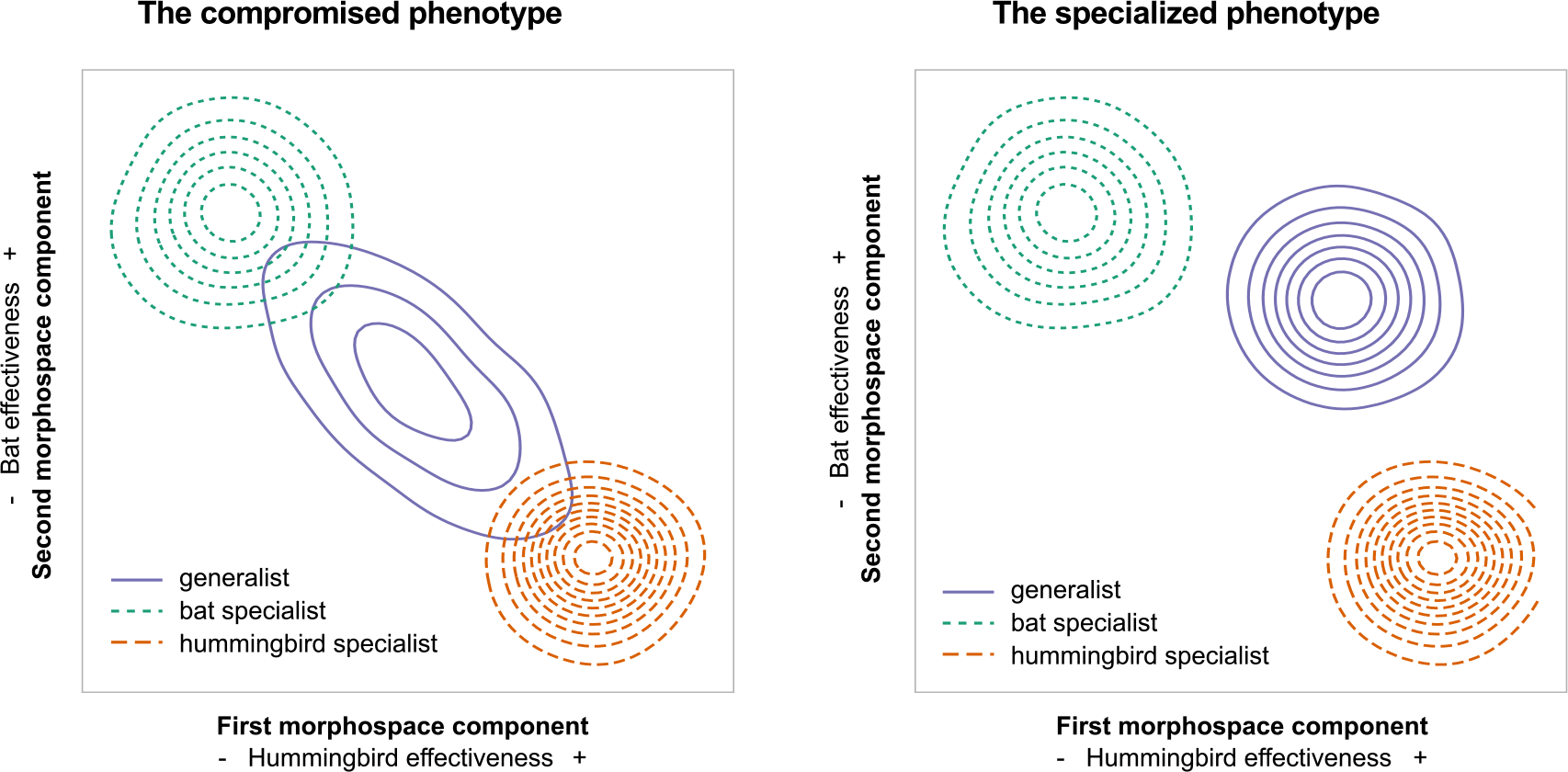
Conceptual phenotypic landscapes illustrating the distribution in mean phenotype of ecological specialist and generalist species associated with two scenarios of evolution of generalization. The x- and y-axis are associated with increases in hummingbird and bat pollination effectiveness, respectively. This is clearly an oversimplified example as real morphospaces are multi-dimensional and the relationship between pollination effectiveness and morphology more complex. In the **compromised phenotype** scenario, the generalists are intermediate in morphology between the two specialists and have the potential to occupy the full extent of variation between the two extremes. In such a scenario, the generalists can have greater variance in morphology and pollination effectiveness than specialists. In the **specialized phenotype** scenario, the generalists occupy a completely distinct region of the morphological landscape; they do not (necessarily) show increased morphological variance compared to specialists and shows good pollination effectiveness by both types of pollinators.

An alternative, called the specialized phenotype, is that the generalists occupy a distinct region in the phenotypic landscape (Fig. 5). Under this scenario, the morphological variance in floral shape need not be greater than that of specialists as generalists could have a floral morphology optimized to both types of pollinators. This model may fit well to the present group as it has been suggested that the presence of a constriction at the base of the corolla for species with a mixed-pollination strategy could represent an adaptation to allow a good pollination service by both hummingbirds and bats by forcing them to approach the flower in a specific way (Martén-Rodríguez et al., 2009). The fact that the corolla shape typical of this pollination strategy has evolved recurrently in the group (Fig. 2) certainly adds weight to this hypothesis. These mixed-pollination species might thus have a phenotypically specialized corolla, in the sense that it is well adapted to both bat and hummingbird pollination, even though they are ecological generalists by being pollinated by different functional pollinators. Indeed, concepts of phenotypic specialization and ecological specialization need not be correlated (Ollerton et al., 2007; Fleming and Muchhala, 2008; Armbruster, 2014). This strategy might be particularly successful in fine-grained pollination environment (Aigner, 2006), such as where pollinators are scarce or vary through time (Waser et al., 1996). Such hypothesis of adaptive generalization (see Gómez and Zamora, 2006) certainly deserves more attention in the future, and will require information on pollination frequency and efficiency to properly associate flower shape to the relative efficiency of pollinators.

The detection of selection contraints for both pollination strategies is noteworthy given that several factors probably contribute in reducing this signal over macroevolutionary timescales. For instance, temporal variation in pollination guilds over macroevolutionary times could weaken the signal of selection, mirroring observations at the population level (e.g., Campbell, 1989; Campbell et al., 1991). The pollination guilds were assumed to be functionally constant over time in our analyses. But given that the exact species pollinating the flowers vary among plant species (Martén-Rodríguez et al., 2009, 2015), the whole story might be more complex. For instance, unrecognized sub-syndromes could be responsible for the larger variation observed for the hummingbird strategy and additional pollinator information will be needed to investigate this further. Variation in selective pressure among species could also occur if agents other than pollinators affect corolla shape. For instance, the apical constriction of the corolla of hummingbird pollinated *Drymonia* (Gesneriaceae) has recently been suggested to be an adaptation to exclude bees (Clark et al., 2015). Moreover, herbivores, including nectar robbers, may affect the selective forces imposed on flowers by pollinators (e.g., Galen and Cuba, 2001; Gómez, 2003). While non-pollinating floral visitors-including bees-are generally not abundant in the group (Martén-Rodríguez et al., 2009, 2015) and herbivory is not common (pers. obs.), it is difficult to completely discard this possibility.

This study is one of the first to show evidence of constrained evolution on flower shapes imposed by pollinator guilds over macroevolutionary time scales and demonstrates the usefulness of a phylogenetic approach to understand pollinator mediated selection. Although additional investigations are needed to confirm these patterns, this study certainly adds weight to the evidence accumulated by many others over the years that the specialist - generalist continuum in pollination biology is complex (Waser et al., 1996; Waser and Ollerton, 2006) and that we cannot assume a priori that pollination specialists show reduced phenotypic disparity compared to pollination generalists.

## Supplementary figures and tables

The supplementary figures and table are available with the supplementary material that is associated with this manuscript. The interactive supplementary figures can be visualized here: www.plantevolution.org/data/Joly_2017_SuppFigs.html.

## Authors contributions

SJ conceived the study. FL, HA, ELB and JLC collected the data, SJ, FL)and JC analyzed the data, SJ wrote the draft and all authors contributed and critically edited the final manuscript.

## Acknowledgements

We thank William Cinea and Phito Merizier that significantly contributed to making this work possible, Julie Faure for constructive discussions, and Cécile Ané for comments and suggestions on a previous manuscript. We also acknowledge the help of Calcul Québec and the Genome Québec Innovation Centre.

---

**Table S1:**
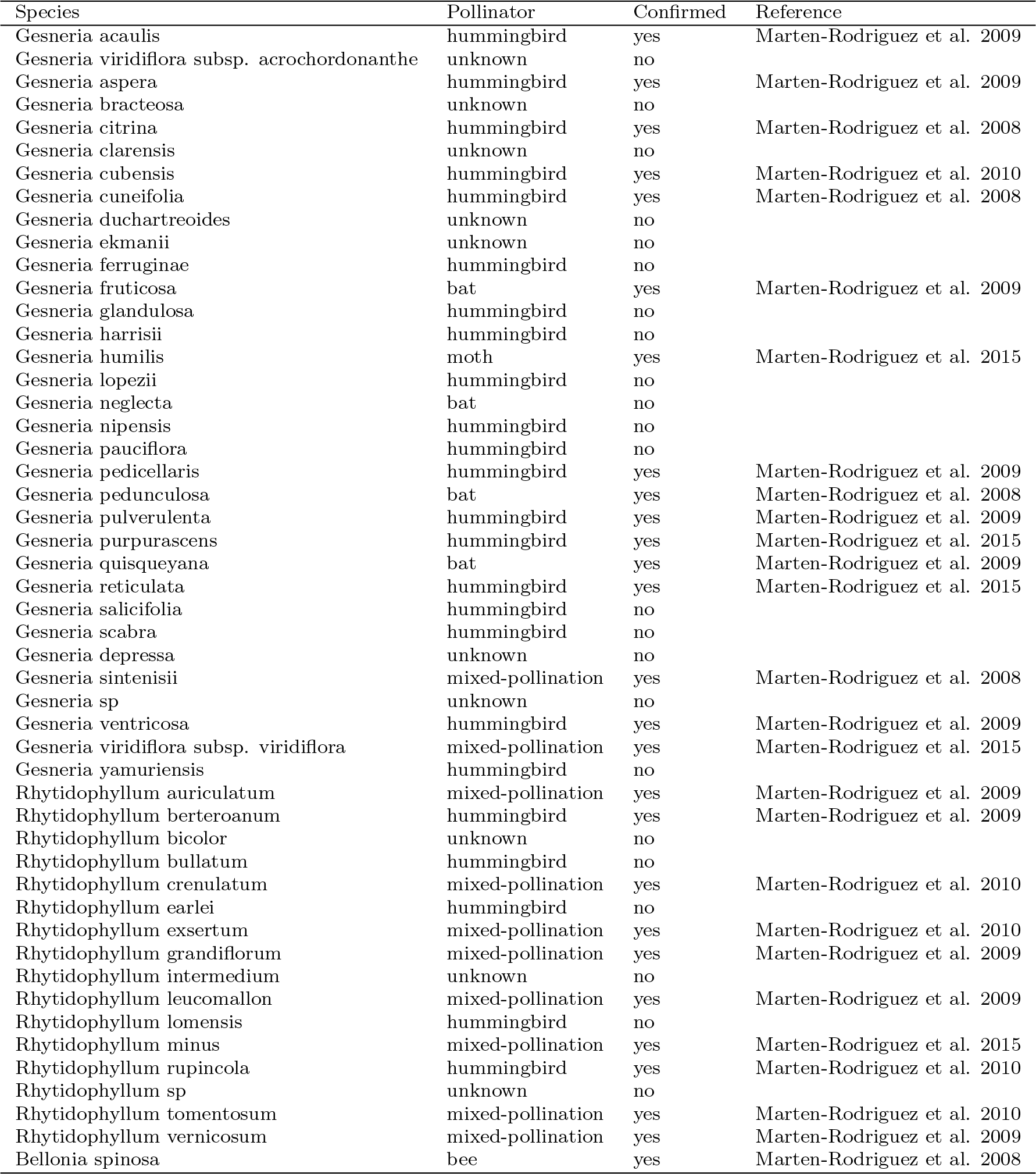
Pollinator information for the species included in the study.

| Species | Pollinator | Confirmed | Reference |
| --- | --- | --- | --- |
| *Gesneria acaulis* | hummingbird | yes | Marten-Rodriguez et al. 2009 |
| *Gesneria viridiflora* subsp. *acrochordonanthe* | unknown | no |  |
| *Gesneria aspera* | hummingbird | yes | Marten-Rodriguez et al. 2009 |
| *Gesneria bracteosa* | unknown | no |  |
| *Gesneria citrina* | hummingbird | yes | Marten-Rodriguez et al. 2008 |
| *Gesneria clarensis* | unknown | no |  |
| *Gesneria cubensis* | hummingbird | yes | Marten-Rodriguez et al. 2010 |
| *Gesneria cuneifolia* | hummingbird | yes | Marten-Rodriguez et al. 2008 |
| *Gesneria duchartreoides* | unknown | no |  |
| *Gesneria ekmanii* | unknown | no |  |
| *Gesneria ferruginae* | hummingbird | no |  |
| *Gesneria fruticosa* | bat | yes | Marten-Rodriguez et al. 2009 |
| *Gesneria glandulosa* | hummingbird | no |  |
| *Gesneria harrisii* | hummingbird | no |  |
| *Gesneria humilis* | moth | yes | Marten-Rodriguez et al. 2015 |
| *Gesneria lopezii* | hummingbird | no |  |
| *Gesneria neglecta* | bat | no |  |
| *Gesneria nipensis* | hummingbird | no |  |
| *Gesneria pauciflora* | hummingbird | no |  |
| *Gesneria pedicellaris* | hummingbird | yes | Marten-Rodriguez et al. 2009 |
| *Gesneria pedunculosa* | bat | yes | Marten-Rodriguez et al. 2008 |
| *Gesneria pulverulenta* | hummingbird | yes | Marten-Rodriguez et al. 2009 |
| *Gesneria purpurascens* | hummingbird | yes | Marten-Rodriguez et al. 2015 |
| *Gesneria quisqueyana* | bat | yes | Marten-Rodriguez et al. 2009 |
| *Gesneria reticulata* | hummingbird | yes | Marten-Rodriguez et al. 2015 |
| *Gesneria salicifolia* | hummingbird | no |  |
| *Gesneria scabra* | hummingbird | no |  |
| *Gesneria depressa* | unknown | no |  |
| *Gesneria sintenisii* | mixed-pollination | yes | Marten-Rodriguez et al. 2008 |
| *Gesneria sp* | unknown | no |  |
| *Gesneria ventricosa* | hummingbird | yes | Marten-Rodriguez et al. 2009 |
| *Gesneria viridiflora* subsp. *viridiflora* | mixed-pollination | yes | Marten-Rodriguez et al. 2015 |
| *Gesneria yamuriensis* | hummingbird | no |  |
| *Rhytidophyllum auriculatum* | mixed-pollination | yes | Marten-Rodriguez et al. 2009 |
| *Rhytidophyllum berterioanum* | hummingbird | yes | Marten-Rodriguez et al. 2009 |
| *Rhytidophyllum bicolor* | unknown | no |  |
| *Rhytidophyllum bullatum* | hummingbird | no |  |
| *Rhytidophyllum crenulatum* | mixed-pollination | yes | Marten-Rodriguez et al. 2010 |
| *Rhytidophyllum earlei* | hummingbird | no |  |
| *Rhytidophyllum exsertum* | mixed-pollination | yes | Marten-Rodriguez et al. 2010 |
| *Rhytidophyllum grandiflorum* | mixed-pollination | yes | Marten-Rodriguez et al. 2009 |
| *Rhytidophyllum intermedium* | unknown | no |  |
| *Rhytidophyllum leucomallum* | mixed-pollination | yes | Marten-Rodriguez et al. 2009 |
| *Rhytidophyllum lomensis* | hummingbird | no |  |
| *Rhytidophyllum minus* | mixed-pollination | yes | Marten-Rodriguez et al. 2015 |
| *Rhytidophyllum rupicola* | hummingbird | yes | Marten-Rodriguez et al. 2010 |
| *Rhytidophyllum sp* | unknown | no |  |
| *Rhytidophyllum tomentosum* | mixed-pollination | yes | Marten-Rodriguez et al. 2010 |
| *Rhytidophyllum vernicosum* | mixed-pollination | yes | Marten-Rodriguez et al. 2009 |
| *Bellonia spinosa* | bee | yes | Marten-Rodriguez et al. 2008 |

**Table S2:**
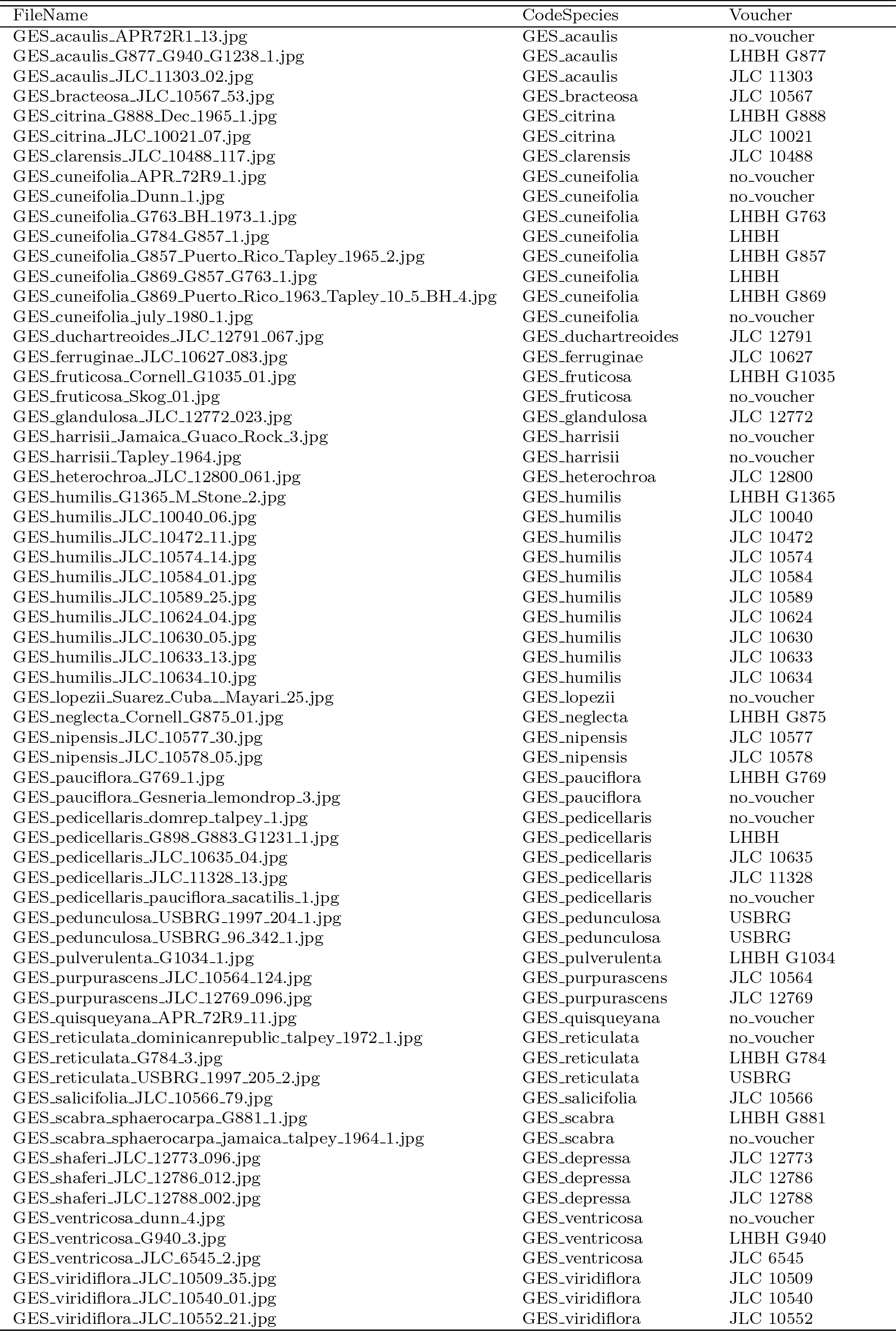
Information on the flower pictures included in the study.

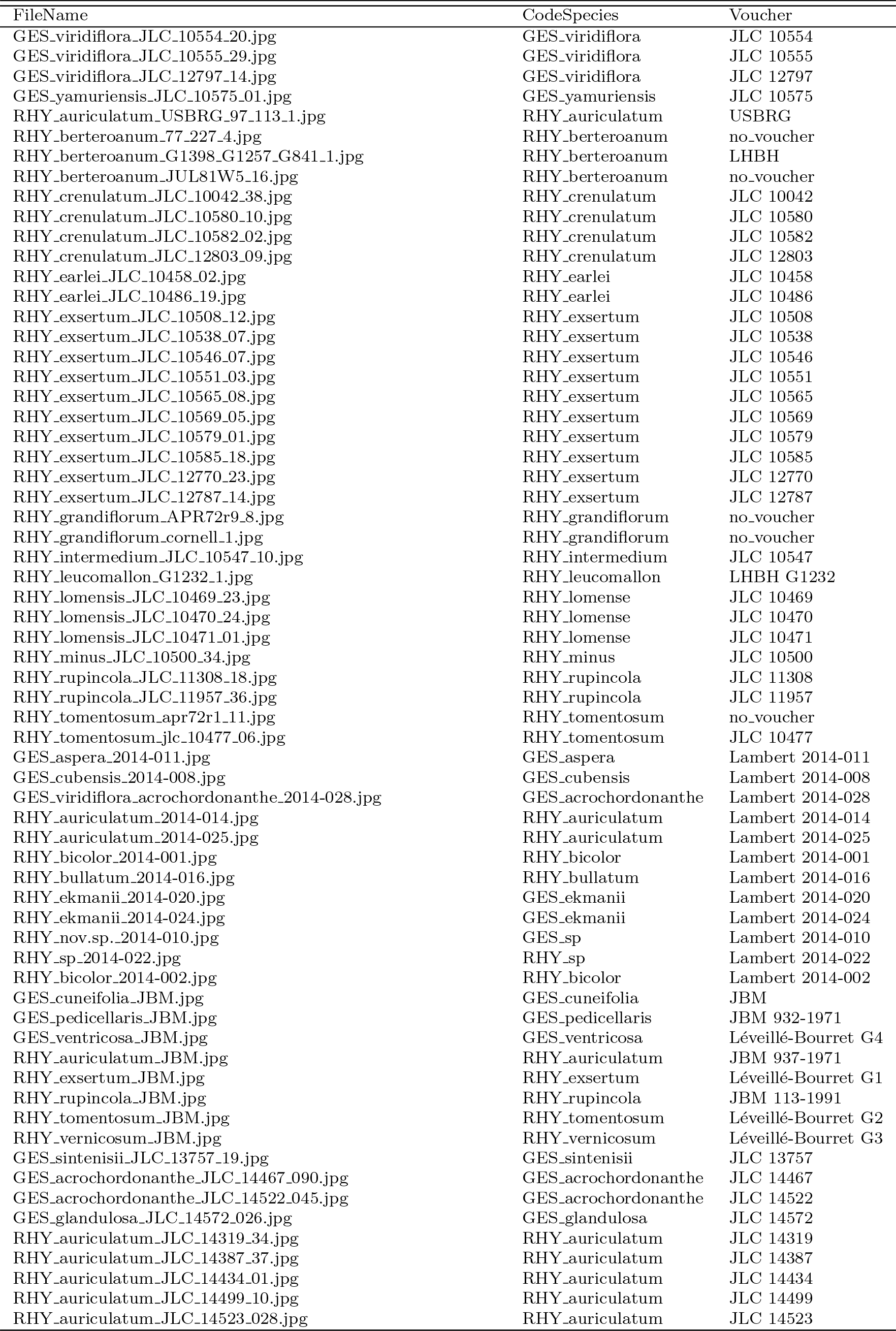

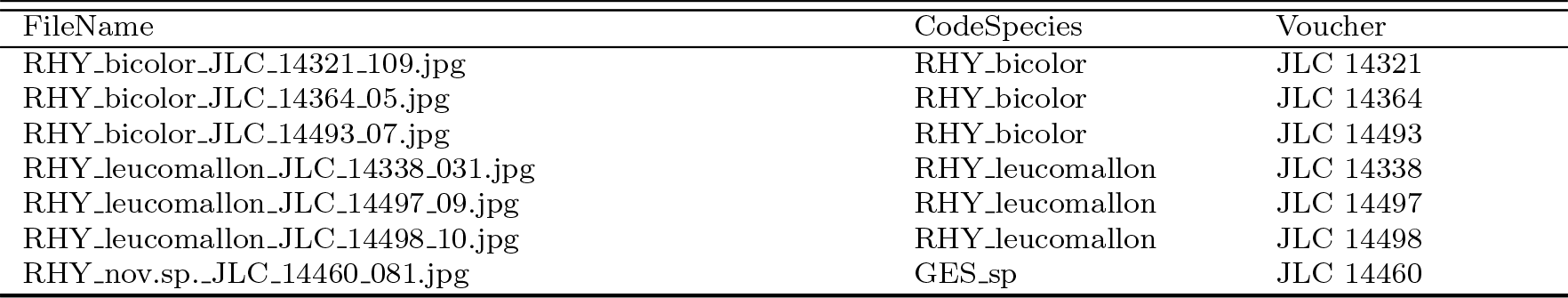

**Table S3:** Voucher information for the specimens sequenced in the study.

| Species | Collector | Collection number | Herbarium | Museum ID | F3H | uf3gt | chi | cycloidea | gapdh |
| --- | --- | --- | --- | --- | --- | --- | --- | --- | --- |
| Bellonia spinosa | Clark, J. | 10573 | UNA | MT-179392 |  |  | sequenced | sequenced | sequenced |
| Bellonia spinosa | Léveillé-Bourret, É. | G8 | MT | MT-184105 | sequenced | sequenced | sequenced |  |  |
| Gesneria acaulis | Clark, J. | 11303 | UNA | MT-179400 | sequenced |  | sequenced | sequenced | sequenced |
| Gesneria acaulis | Martén-Rodriguez, S. | US-3479058-1188 | US |  |  |  |  | GU323229 |  |
| Gesneria acrochordonanthe | Lambert, F. | 2014-027 | MT | MT-194054 |  |  | sequenced | sequenced |  |
| Gesneria acrochordonanthe | Lambert, F. | 2014-028 | MT | MT-194056 | sequenced | sequenced | sequenced | sequenced |  |
| Gesneria aspera | Lambert, F. | 2014-011 | MT | MT-194032 | sequenced |  | sequenced | sequenced |  |
| Gesneria bracteosa | Clark, J. | 10567 | UNA | MT-179575 | sequenced | sequenced | sequenced | sequenced | sequenced |
| Gesneria christii | Clark, J. | 10025 | UNA |  | sequenced |  | sequenced | sequenced |  |
| Gesneria christii | Léveillé-Bourret, É. | G6 | MT | MT-184103 | sequenced | sequenced | sequenced | sequenced | sequenced |
| Gesneria christii | ?? | SI 94-507 | ?? |  |  |  |  | AY363923 |  |
| Gesneria citrina | Clark, J. | 10020 | UNA |  | sequenced |  |  | sequenced |  |
| Gesneria citrina | Martén-Rodriguez, S. | 1248 |  |  |  |  |  | GU323232 |  |
| Gesneria clarensis | Clark, J. | 10488 | UNA | MT-179401 | sequenced | sequenced | sequenced | sequenced | sequenced |
| Gesneria cubensis | Lambert, F. | 2014-008 | MT | MT-194027 | sequenced |  | sequenced | sequenced |  |
| Gesneria cubensis | Martén-Rodriguez, S. | JBSD-109650-1232 | UNA |  |  |  |  | GU323234 |  |
| Gesneria cuneifolia | Martén-Rodriguez, S. | 1247 |  |  |  |  |  | GU323235 |  |
| Gesneria ekmanii | Lambert, F. | 2014-018 | MT | MT-194044 | sequenced | sequenced | sequenced | sequenced |  |
| Gesneria ekmanii | Lambert, F. | 2014-020 | MT | MT-194046 | sequenced | sequenced | sequenced | sequenced | sequenced |
| Gesneria ekmanii | Lambert, F. | 2014-024 | MT | MT-194050 | sequenced |  | sequenced | sequenced |  |
| Gesneria ferruginea | Clark, J. | 10627 | UNA | MT-179294 |  | sequenced | sequenced | sequenced | sequenced |
| Gesneria fruticosa | Lambert, F. | 2014-012 | MT | MT-194035 | sequenced | sequenced | sequenced | sequenced | sequenced |
| Gesneria fruticosa | Martén-Rodriguez, S. | JBSD-109606-1227 | UNA | MT-179391 |  |  |  | GU323238 |  |
| Gesneria humilis | Clark, J. | 10626 | UNA | MT-179385 | sequenced | sequenced | sequenced | sequenced | sequenced |
| Gesneria humilis | Clark, J. | 10040 | UNA | MT-179363 | sequenced | sequenced | sequenced | sequenced | sequenced |
| Gesneria humilis | Clark, J. | 10472 | UNA |  | sequenced | sequenced | sequenced | sequenced | sequenced |
| Gesneria humilis | Clark, J. | 10574 | UNA |  |  |  | sequenced | sequenced |  |
| Gesneria humilis | Chautems | 1179 |  |  |  |  |  | AY423156 |  |
| Gesneria nipensis | Clark, J. | 10577 | UNA | MT-179356 | sequenced |  |  | sequenced |  |
| Gesneria pedicellaris | Clark, J. | 10635 | UNA | MT-179354 | sequenced | sequenced | sequenced | sequenced |  |
| Gesneria pedicellaris | Martén-Rodriguez, S. | US-3498534-1229 | US |  |  |  |  | GU323241 |  |
| Gesneria pedunculosa | Martén-Rodriguez, S. | 1251 |  |  |  |  |  | GU323242 |  |
| Gesneria pedunculosa | Clark, J. | 10644 | UNA | MT-179353 | sequenced | sequenced | sequenced | sequenced |  |
| Gesneria pulverulenta | Martén-Rodriguez, S. | US-3498527-1237 | US |  |  |  |  | GU323243 |  |
| Gesneria pumila | Martén-Rodriguez, S. | 1194 | UNA | MT-179573 |  |  |  | GU323244 |  |
| Gesneria purpurascens | Clark, J. | 10564 | UNA | MT-179351 | sequenced | sequenced | sequenced | sequenced | sequenced |
| Gesneria quisqueyana | Martén-Rodriguez, S. | US-3498533-1230 | US |  |  |  |  | GU323245 |  |
| Gesneria reticulata | Clark, J. | 10558 | UNA | MT-179347 | sequenced | sequenced | sequenced | sequenced |  |
| Gesneria reticulata | Martén-Rodriguez, S. | US-3498539-1221 | US |  |  |  |  | GU323246 |  |
| Gesneria salicifolia | Clark, J. | 10566 | UNA | MT-179316 | sequenced |  | sequenced | sequenced | sequenced |
| Gesneria depressa | Clark, J. | 13070 | UNA |  | sequenced |  |  | sequenced |  |
| Gesneria sintenisii | Clark, J. | 13757 | UNA |  | sequenced | sequenced | sequenced | sequenced |  |
| Gesneria sintenisii | Martén-Rodriguez, S. | 1252 |  | US-3594111 |  |  |  | GU323250 |  |
| Gesneria ventricosa | Clark, J. | 6545 | UNA | MT-179344 | sequenced |  |  | sequenced | sequenced |
| Gesneria ventricosa | Léveillé-Bourret, É. | G4 | MT | MT-184101 | sequenced | sequenced | sequenced | sequenced | sequenced |
| Gesneria ventricosa | Martén-Rodriguez, S. | 1112A |  |  |  |  |  | GU323249 |  |

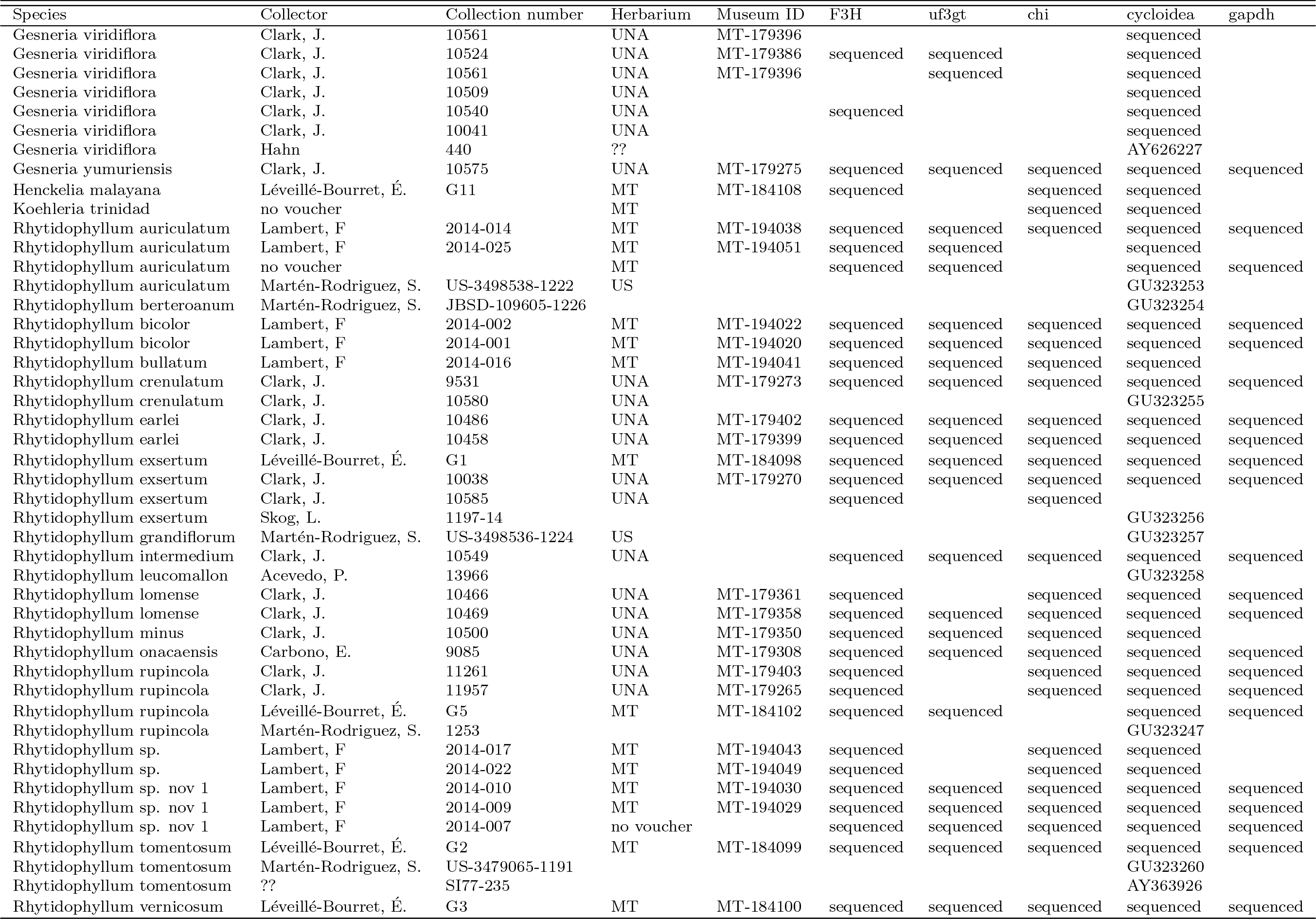

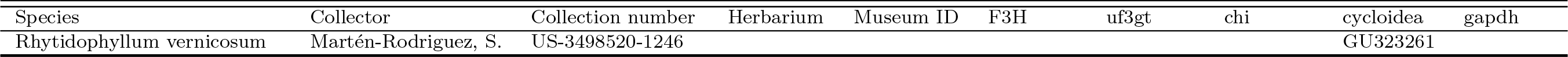

**Table S4:**
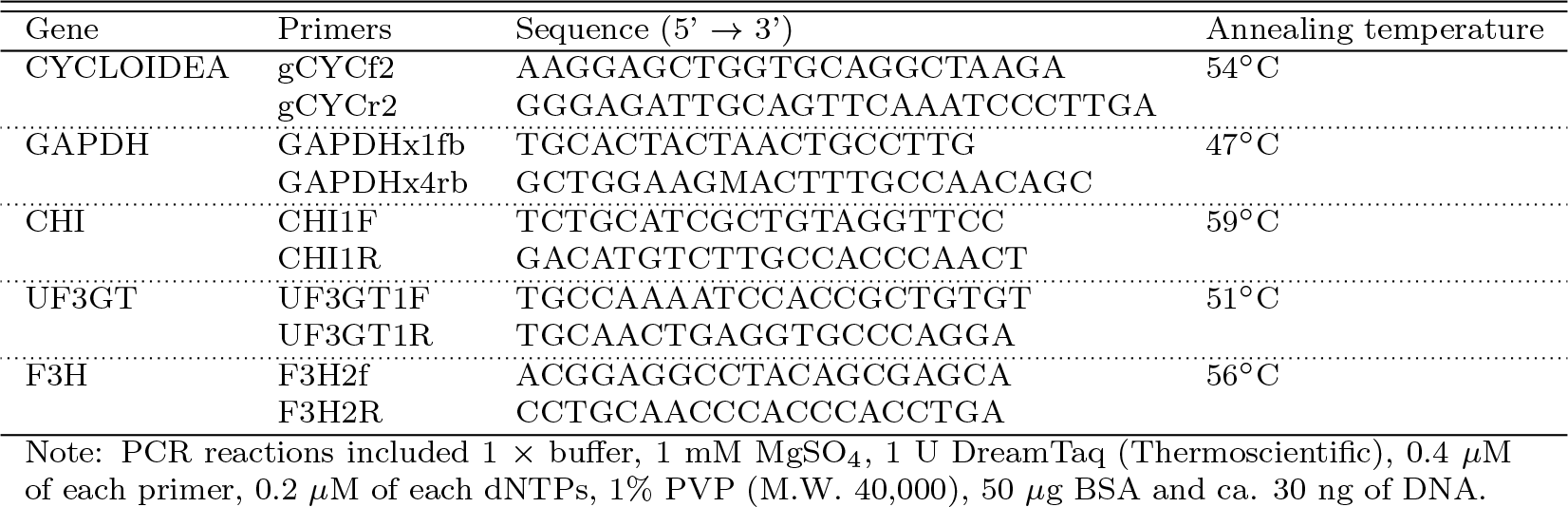
Primer information for the gene amplification.

**Table S5:** Number of transitions between the different pollination strategies according to the stochastic mapping when performed on species with confirmed and inferred pollination strategies. The median values obtained from the character simulations over the posterior distribution of species tree is reported as well as 95% credible intervals. Ancestral state are in rows

|  | bat | bee | hummingbird | mixed-pollination | moth |
| --- | --- | --- | --- | --- | --- |
| bat | – | 0.25 [0.19, 0.22] | 3.74 [3.07, 4.40] | 3.70 [3.06, 4.30] | 0.22 [0.14, 0.34] |
| bee | 0.06 [0.02, 0.10] | – | 0.05 [0.02, 0.10] | 0.08 [0.02, 0.14] | 0.03 [0.02, 0.05] |
| hummingbird | 3.32 [2.55, 3.88] | 0.43 [0.37, 0.51] | – | 2.48 [2.00, 2.96] | 0.63 [0.44, 0.78] |
| mixed-pollination | 4.47 [3.82, 5.17] | 0.28 [0.20, 0.37] | 6.53 [5.54, 7.15] | – | 0.25 [0.14, 0.33] |
| moth | 0.03 [0.02, 0.11] | 0.06 [0.04, 0.10] | 0.04 [0.01, 0.07] | 0.04 [0.02, 0.07] | – |

**Table S6:** Parameter values of the univariate evolutionary models fitted on the first three principal components of the morphospace when species with confirmed and inferred pollinators were included in the analyses. Mean values from the posterior distribution of species trees are given for the AICc weights, whereas median values are given for the parameter estimates. Numbers in brackets indicate the 25% and the 75% quantiles. The best model for each component is in bold. The *θ* parameter indicate the global or regime means for the BM-type and OUBM-type models, whereas it indicates the stationary optimum trait for the OU-type models, *station*_*hum*_ and *station*_*mix*_ are the stationary distributions of the hummingbird and mixed-pollination strategies.

| PC1 |  |  |  |  |  |  |  |  |  |  |
| --- | --- | --- | --- | --- | --- | --- | --- | --- | --- | --- |
| Models | p | AICc weight | $\theta_{hum}$ | $\theta_{mix}$ | $\sigma_{hum}^2$ | $\sigma_{mix}^2$ | $station_{hum}$ | $station_{mix}$ | | |
| BM1 | 2 | 0 [0,0] | 0.042 [0.033,0.05] | 0.042 [0.033,0.05] | 0.105 [0.077,0.163] | 0.105 [0.077,0.163] | — | — |  |  |
| BMV | 3 | 0 [0,0] | 0.097 [0.033,0.137] | 0.097 [0.033,0.137] | 0.028 [0.016,0.13] | 0.151 [0.059,0.285] | — | — |  |  |
| BM1m | 3 | 0.38 [0.03,0.69] | 0.154 [0.148,0.162] | -0.129 [-0.139,-0.118] | 0.015 [0.012,0.019] | 0.015 [0.012,0.019] | — | — |  |  |
| BMVm | 4 | 0.16 [0.03,0.24] | 0.153 [0.143,0.16] | -0.132 [-0.142,-0.122] | 0.019 [0.013,0.025] | 0.011 [0.009,0.014] | — | — |  |  |
| OU1 | 3 | 0 [0,0] | 0.031 [0.021,0.042] | 0.031 [0.021,0.042] | 0.207 [0.137,0.477] | 0.207 [0.137,0.477] | 0.033 [0.031,0.034] | 0.033 [0.031,0.034] |  |  |
| OUM | 4 | 0.46 [0.03,0.95] | 0.169 [0.165,0.172] | -0.159 [-0.159,-0.158] | 3.122 [0.916,17.092] | 3.122 [0.916,17.092] | 0.005 [0.005,0.005] | 0.005 [0.005,0.005] |  |  |
| OUMV | 5 | 0 [0,0] | 0.214 [0.192,0.23] | -0.163 [-0.192,-0.075] | 15.291 [4.636,43.424] | 24.445 [10.766,50.789] | 2.25 [1.027,5.547] | 4.309 [1.556,7.299] |  |  |
| OUMA | 5 | 0 [0,0] | 0.218 [0.198,0.247] | -0.159 [-0.196,-0.08] | 21.291 [12.123,39.902] | 21.291 [12.123,39.902] | 3.587 [2.64,4.543] | 3.587 [2.64,4.543] |  |  |
| OUMVA | 6 | 0 [0,0] | 0.214 [0.19,0.231] | -0.162 [-0.195,-0.096] | 17.889 [4.882,42.186] | 24.258 [10.321,52.843] | 2.414 [1.005,4.92] | 4.403 [1.848,7.136] |  |  |
| OUBMi | 4 | 0 [0,0] | 0.123 [0.096,0.146] | 0.123 [0.096,0.146] | 1.797 [0.063,43.782] | 0.038 [0.018,0.119] | 0.014 [0.007,0.017] | — |  |  |
| BMOUi | 4 | 0 [0,0] | -0.053 [-0.074,-0.03] | -0.053 [-0.074,-0.03] | 0.033 [0.021,0.066] | 2.751 [0.442,33.744] | — | 0.019 [0.013,0.023] |  |  |
| OUBM | 3 | 0 [0,0] | 0.15 [0.118,0.177] | 0.15 [0.118,0.177] | 0.171 [0.118,0.285] | 0.171 [0.118,0.285] | 0.005 [0.004,0.007] | — |  |  |
| BMOU | 3 | 0 [0,0] | -0.091 [-0.119,-0.036] | -0.091 [-0.119,-0.036] | 0.188 [0.122,0.336] | 0.188 [0.122,0.336] | — | 0.008 [0.006,0.011] |  |  |
| PC2 |  |  |  |  |  |  |  |  |  |  |
| Models | p | AICc weight | $\theta_{hum}$ | $\theta_{mix}$ | $\sigma_{hum}^2$ | $\sigma_{mix}^2$ | $station_{hum}$ | $station_{mix}$ | | |
| BM1 | 2 | 0 [0,0] | -0.033 [-0.036,-0.03] | -0.033 [-0.036,-0.03] | 0.021 [0.016,0.035] | 0.021 [0.016,0.035] | — | — |  |  |
| BMV | 3 | 0 [0,0] | -0.032 [-0.037,-0.027] | -0.032 [-0.037,-0.027] | 0.029 [0.02,0.055] | 0.009 [0.007,0.012] | — | — |  |  |
| BM1m | 3 | 0 [0,0] | -0.018 [-0.024,-0.012] | -0.057 [-0.064,-0.051] | 0.019 [0.014,0.033] | 0.019 [0.014,0.033] | — | — |  |  |
| BMVm | 4 | 0 [0,0] | -0.012 [-0.02,0] | -0.05 [-0.059,-0.039] | 0.028 [0.019,0.052] | 0.007 [0.005,0.009] | — | — |  |  |
| OU1 | 3 | 0.53 [0.45,0.65] | -0.028 [-0.03,-0.026] | -0.028 [-0.03,-0.026] | 0.223 [0.075,0.99] | 0.223 [0.075,0.99] | 0.003 [0.003,0.003] | 0.003 [0.003,0.003] |  |  |
| OUM | 4 | 0.26 [0.23,0.32] | -0.04 [-0.043,-0.034] | -0.017 [-0.019,-0.014] | 0.169 [0.075,0.662] | 0.169 [0.075,0.662] | 0.003 [0.003,0.003] | 0.003 [0.003,0.003] |  |  |
| OUMV | 5 | 0.05 [0.01,0.07] | -0.049 [-0.055,-0.038] | -0.02 [-0.027,-0.012] | 9.117 [5.89,13.777] | 3.756 [2.774,5.29] | 1.132 [0.884,1.613] | 0.482 [0.408,0.568] |  |  |
| OUMA | 5 | 0.02 [0.0,0.03] | -0.049 [-0.056,-0.039] | -0.016 [-0.026,-0.008] | 6.422 [4.583,9.904] | 6.422 [4.583,9.904] | 0.796 [0.7,0.974] | 0.796 [0.7,0.974] |  |  |
| OUMVA | 6 | 0.01 [0.0,0.02] | -0.049 [-0.056,-0.038] | -0.02 [-0.028,-0.012] | 9.189 [5.72,13.364] | 3.717 [2.825,5.463] | 1.117 [0.865,1.458] | 0.486 [0.411,0.583] |  |  |
| OUBMi | 4 | 0.1 [0.01,0.06] | -0.041 [-0.047,-0.035] | -0.041 [-0.047,-0.035] | 0.289 [0.11,1.407] | 0.006 [0.004,0.009] | 0.003 [0.003,0.003] | — |  |  |
| BMOUi | 4 | 0.01 [0,0] | -0.023 [-0.033,-0.017] | -0.023 [-0.033,-0.017] | 0.028 [0.019,0.049] | 0.045 [0.023,0.225] | — | 0.002 [0.002,0.002] |  |  |
| OUBM | 3 | 0 [0,0] | -0.045 [-0.051,-0.037] | -0.045 [-0.051,-0.037] | 0.032 [0.021,0.05] | 0.032 [0.021,0.05] | 0.003 [0.003,0.004] | — |  |  |
| BMOU | 3 | 0 [0,0] | -0.024 [-0.033,-0.017] | -0.024 [-0.033,-0.017] | 0.03 [0.021,0.051] | 0.03 [0.021,0.051] | — | 0.002 [0.002,0.002] |  |  |
| PC3 |  |  |  |  |  |  |  |  |  |  |
| Models | p | AICc weight | $\theta_{hum}$ | $\theta_{mix}$ | $\sigma_{hum}^2$ | $\sigma_{mix}^2$ | $station_{hum}$ | $station_{mix}$ | | |
| BM1 | 2 | 0.02 [0.0,0.03] | 0.018 [0.015,0.019] | 0.018 [0.015,0.019] | 0.004 [0.004,0.005] | 0.004 [0.004,0.005] | — | — |  |  |
| BMV | 3 | 0.01 [0,0.01] | 0.017 [0.015,0.019] | 0.017 [0.015,0.019] | 0.005 [0.005,0.007] | 0.003 [0.003,0.005] | — | — |  |  |
| BM1m | 3 | 0.12 [0.03,0.16] | 0.031 [0.028,0.034] | -0.004 [-0.009,-0.001] | 0.004 [0.003,0.004] | 0.004 [0.003,0.004] | — | — |  |  |
| BMVm | 4 | 0.05 [0.02,0.08] | 0.035 [0.03,0.039] | -0.001 [-0.004,0.003] | 0.005 [0.004,0.006] | 0.002 [0.002,0.003] | — | — |  |  |
| OU1 | 3 | 0.3 [0.13,0.45] | 0.015 [0.014,0.016] | 0.015 [0.014,0.016] | 0.019 [0.013,0.107] | 0.019 [0.013,0.107] | 0.002 [0.002,0.002] | 0.002 [0.002,0.002] |  |  |
| OUM | 4 | 0.1 [0.03,0.15] | 0.018 [0.016,0.021] | 0.01 [0.009,0.013] | 0.02 [0.013,0.209] | 0.02 [0.013,0.209] | 0.002 [0.001,0.002] | 0.002 [0.001,0.002] |  |  |
| OUMV | 5 | 0.12 [0.03,0.16] | 0.022 [0.017,0.029] | 0.01 [0.005,0.017] | 2.865 [2.249,3.919] | 0.854 [0.702,1.063] | 0.449 [0.394,0.524] | 0.146 [0.118,0.17] |  |  |
| OUMA | 5 | 0.03 [0.01,0.04] | 0.022 [0.016,0.026] | 0.01 [0.005,0.018] | 1.359 [1.089,1.744] | 1.359 [1.089,1.744] | 0.275 [0.249,0.301] | 0.275 [0.249,0.301] |  |  |
| OUMVA | 6 | 0.02 [0.01,0.03] | 0.023 [0.019,0.029] | 0.01 [0.006,0.017] | 2.135 [1.616,2.726] | 0.733 [0.569,0.944] | 0.399 [0.348,0.467] | 0.148 [0.118,0.177] |  |  |
| OUBMi | 4 | 0.01 [0,0.01] | 0.017 [0.014,0.02] | 0.017 [0.014,0.02] | 0.015 [0.01,0.027] | 0.003 [0.002,0.004] | 0.002 [0.002,0.003] | — |  |  |
| BMOUi | 4 | 0.13 [0.02,0.16] | 0.016 [0.014,0.017] | 0.016 [0.014,0.017] | 0.004 [0.004,0.005] | 0.048 [0.02,0.229] | — | 0.001 [0.001,0.001] |  |  |
| OUBM | 3 | 0.01 [0,0.01] | 0.018 [0.015,0.021] | 0.018 [0.015,0.021] | 0.005 [0.004,0.007] | 0.005 [0.004,0.007] | 0.003 [0.002,0.007] | — |  |  |
| BMOU | 3 | 0.08 [0.01,0.1] | 0.015 [0.012,0.017] | 0.015 [0.012,0.017] | 0.006 [0.006,0.008] | 0.006 [0.006,0.008] | — | 0.001 [0.001,0.001] |  |  |

**Table S7:** Matrix of stationary variance estimates obtained with the OUM multivariate model, averaged over the posterior distribution of species trees with only species with confirmed pollination strategies. Median values are reported and numbers in brackets indicate the 25% and the 75% quantiles.

|  | PC1 | PC2 | PC3 |
| --- | --- | --- | --- |
| PC1 | 0.0058 [0.0055, 0.0062] |  |  |
| PC2 | 0.00063 [0.00053, 0.00081] | 0.0029 [0.0029, 0.0030] |  |
| PC3 | -0.0010 [-0.0013, -0.00070] | -0.00058 [-0.00064, -0.00053] | 0.0020 [0.0018, 0.0022] |

**Table S8:** Model performance with the multivariate evolutionary models fitted on the first three principal components of the morphospace when all species were included in the analyses, including those with inferred pollinator strategies. The mean values obtained from the posterior distribution of species trees are given; numbers in brackets indicate the 25% and the 75% quantiles. The best model is in bold.

| models | logLik | param | AICc weight |
| --- | --- | --- | --- |
| BM1 | 108.11 [98.93,120.48] | 9 | 0 [0,0] |
| BMV | 119.69 [112.64,128.69] | 15 | 0 [0,0] |
| BM1m | 140.93 [133.11,152.03] | 12 | 0 [0,0.01] |
| BMVm | 147.9 [140.66,156.66] | 18 | 0 [0,0] |
| OU1 | 136.62 [132.59,140.57] | 15 | 0 [0,0] |
| **OUM** | **164.32 [162.02,166.75]** | **18** | **1 [0.99,1]** |
| OUBM | 123.15 [117.04,131.4] | 15 | 0 [0,0] |
| BMOU | 121.65 [115.22,130.89] | 15 | 0 [0,0] |
| OUBMr | 142.66 [140.17,146.98] | 21 | 0 [0,0] |
| BMOUr | 130.75 [123.82,139.6] | 21 | 0 [0,0] |

**Table S9:** Model parameters for the multivariate OUM model, which was the model that received the highest *AICc* weight (Table S8), when all species are included in the analysis. The mean values obtained from the posterior distribution of species trees are given; numbers in brackets indicate the 25% and the 75% quantiles.

| parameters | PC1 | PC2 | PC3 |
| --- | --- | --- | --- |
| $\theta_{hum}$ | 0.173 [0.169,0.177] | -0.042 [-0.052,-0.037] | 0.015 [0.008,0.019] |
| $\theta_{mix}$ | -0.16 [-0.162,-0.158] | -0.017 [-0.018,-0.014] | 0.011 [0.009,0.016] |
| $\sigma^2$ | 3.132 [0.987,10.306] | 0.7 [0.232,2.32] | 0.091 [0.018,0.372] |
| phylogenetic halftime | 0.001 [0,0.003] | 0.013 [0.004,0.039] | 0.122 [0.079,0.21] |
| stationary variance | 0.004 [0.004,0.005] | 0.003 [0,0] | 0.001 [-0.001,0] |

**Figure S1:**
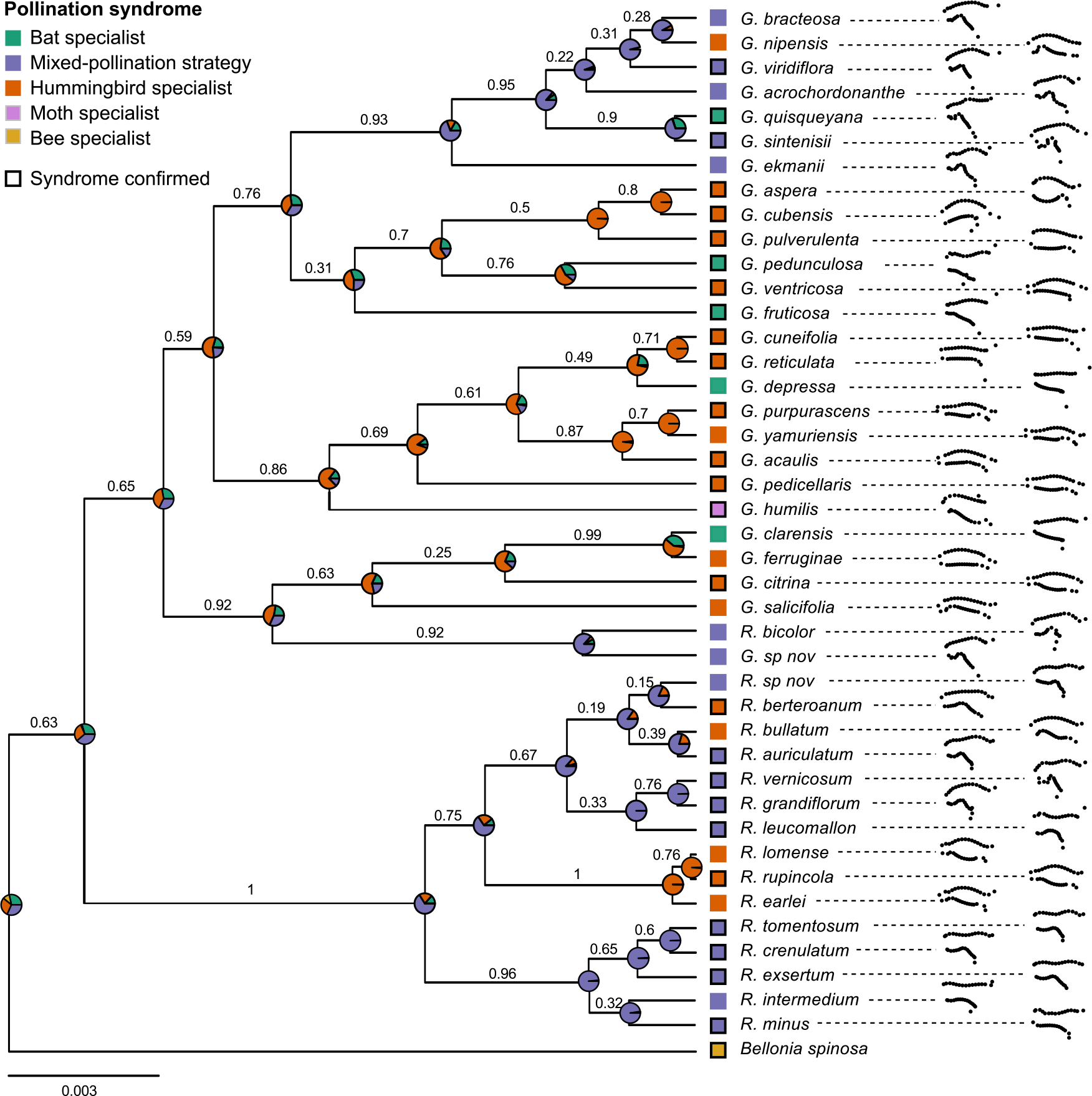
Species phylogeny showing mean corolla shapes (after Procrustes analysis). Pollination strategies are shown with those that have been confirmed indicated by a black contour. Pie charts represent the joint probability of each state at nodes as estimated by stochastic mapping from all species, that is including species with inferred pollinators. Clade posterior probabilities are shown above branches. Outgroup taxa are not shown.

**Figure S2:**
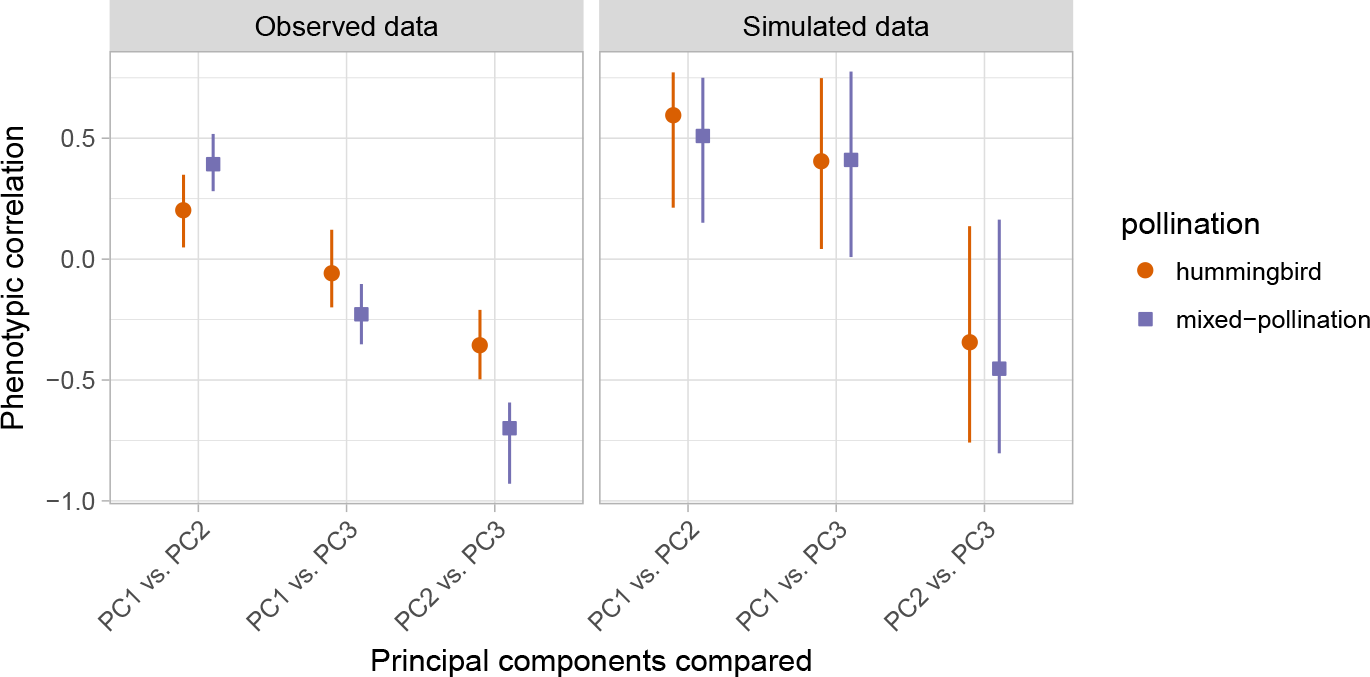
Graphical representation of the evolutionary correlations (i.e., standardized evolutionary rates matrices) obtained with the BMVm multivariate model when all species were included in the analysis, for the observed data (left panel) and for data simulated under the best fitting model (OUM; right panel). Symbols represent the median correlation and the lines the 25% and 75% quantiles for both hummingbirds and mixed-pollination strategies. No artifactual differences are detected between the two groups when fitting models on traits simulated with the OUM model and thus with a common evolutionary covariance (right panel, see text).

